## Supporting Information for "Functional control of a 0.5 MDa TET aminopeptidase by a flexible loop revealed by MAS NMR"

**Table S1.** Kinetic constants derived from fitting the Michaelis–Menten model to the absorbance-increase time traces as TET2 cleaves H-Leu-pNA. See Methods for details.

| TET Mutants | $K_m$ (mM) | $k_{cat}$ ($s^{-1}$) | $k_{cat}/K_m$ ($M^{-1}$ $s^{-1}$) |
|---|---|---|---|
| WT | $3.2 \pm 0.1$ | $14.1 \pm 1.7$ | $4.4 \times 10^3 \pm 0.5 \times 10^3$ |
| H123F | $14.4 \pm 0.6$ | $26.6 \pm 4.1$ | $1.8 \times 10^3 \pm 0.3 \times 10^3$ |
| H123K | $17.0 \pm 1.3$ | $16.9 \pm 3.6$ | $0.9 \times 10^3 \pm 0.2 \times 10^3$ |
| H123Y | $13.8 \pm 0.6$ | $15.7 \pm 2.7$ | $1.1 \times 10^3 \pm 0.2 \times 10^3$ |
| ΔP122,ΔK126 | $28.5 \pm 2.3$ | $31.5 \pm 8.1$ | $1.1 \times 10^3 \pm 0.3 \times 10^3$ |
| P121G,P122G | $43.6 \pm 5.0$ | $55.7 \pm 21.2$ | $1.3 \times 10^3 \pm 0.5 \times 10^3$ |
| Δ-loop (Δ120-138) | $1.3 \pm 0.1$ | $1.7 \pm 1.0$ | $1.3 \times 10^3 \pm 1.0 \times 10^3$ |

**Table S2.** All 29 predicted contacts involving the V120-Q138 loop among the first 2N (706) highest ranked DCA predictions. Whenever these contacts were observed in the MD simulation, the contact type (intra-loop, intramolecular contact within the same subunit or inter-molecular contact) is based on the observed contact frequency in the MD simulations, see Fig. S11. In other cases, the assignment to these different types of contact is tentative, and marked with a "?".

| Contact | Type | Rank |
|---|---|---|
| V120 - D129 | Intra-loop | 81 |
| P89 - W136 | ? | 171 |
| P95 - W136 | ? | 242 |
| K126 - D137 | Intra-molecular | 246 |
| K126 - D135 | Intra-molecular | 272 |
| K96 - A133 | ? | 293 |
| K96 - P134 | ? | 296 |
| I124 - T245 | Inter-molecular | 320 |
| S119 - Q125 | Intra-molecular | 346 |
| I124 - I238 | Inter-molecular | 371 |
| P95 - P134 | ? | 383 |
| P95 - A133 | ? | 399 |
| V120 - Q125 | Intra-loop | 409 |
| D137 - K249 | ? | 416 |
| A118 - H123 | Intra-molecular | 429 |
| P121 - I238 | Inter-molecular | 448 |
| Q125 - R130 | Intra-loop | 464 |
| E80 - Q138 | ? | 487 |
| P127 - A133 | Intra-molecular (?) | 534 |
| I99 - P134 | ? | 539 |
| P95 - D135 | ? | 565 |
| D94 - P122 | Intra-molecular | 568 |
| K132 - Q138 | Intra-molecular | 589 |
| E128 – D135 | Intra-molecular | 603 |
| D94 - D129 | Intra-molecular (?) | 618 |
| V120 - Q138 | Intra-molecular | 629 |
| D94 - Q125 | Intra-molecular (?) | 640 |
| D129 - P134 | Intra-molecular (?) | 679 |
| E128 - P134 | Intra-molecular (?) | 685 |

**Table S3.** Spin-lock radio-frequency fields and durations for the measurement of the $^{13}$C $R_{1\rho}$ experiment.

| radio-frequency (RF) field strengths (kHz) | Relaxation delays (ms) |
|---|---|
| 10.0 | 1, 25, 75, 100, 150, 200, 250 |
| 20.0 | 1, 5, 10, 20, 25, 30,40,55 |
| 30.0 | 1, 5, 10, 15, 20, 30,45,60,100 |
| 33.0 | 1, 5, 10, 15, 20, 25,30,50,80 |
| 35.0 | 1, 5, 10, 15, 20, 25,30,50,80 |
| 38.0 | 1, 5, 10, 15, 20, 25,30,50,80 |
| 41.0 | 1, 5, 7.5, 10, 15, 20,25,30,50 |
| 42.0 | 1, 5, 7.5, 10, 15, 20,25,30,50 |
| 43.0 | 1, 5, 7.5, 10, 15, 20,25,30,50 |
| 44.0 | 1, 5, 7.5, 10, 15, 20,25,30,40 |

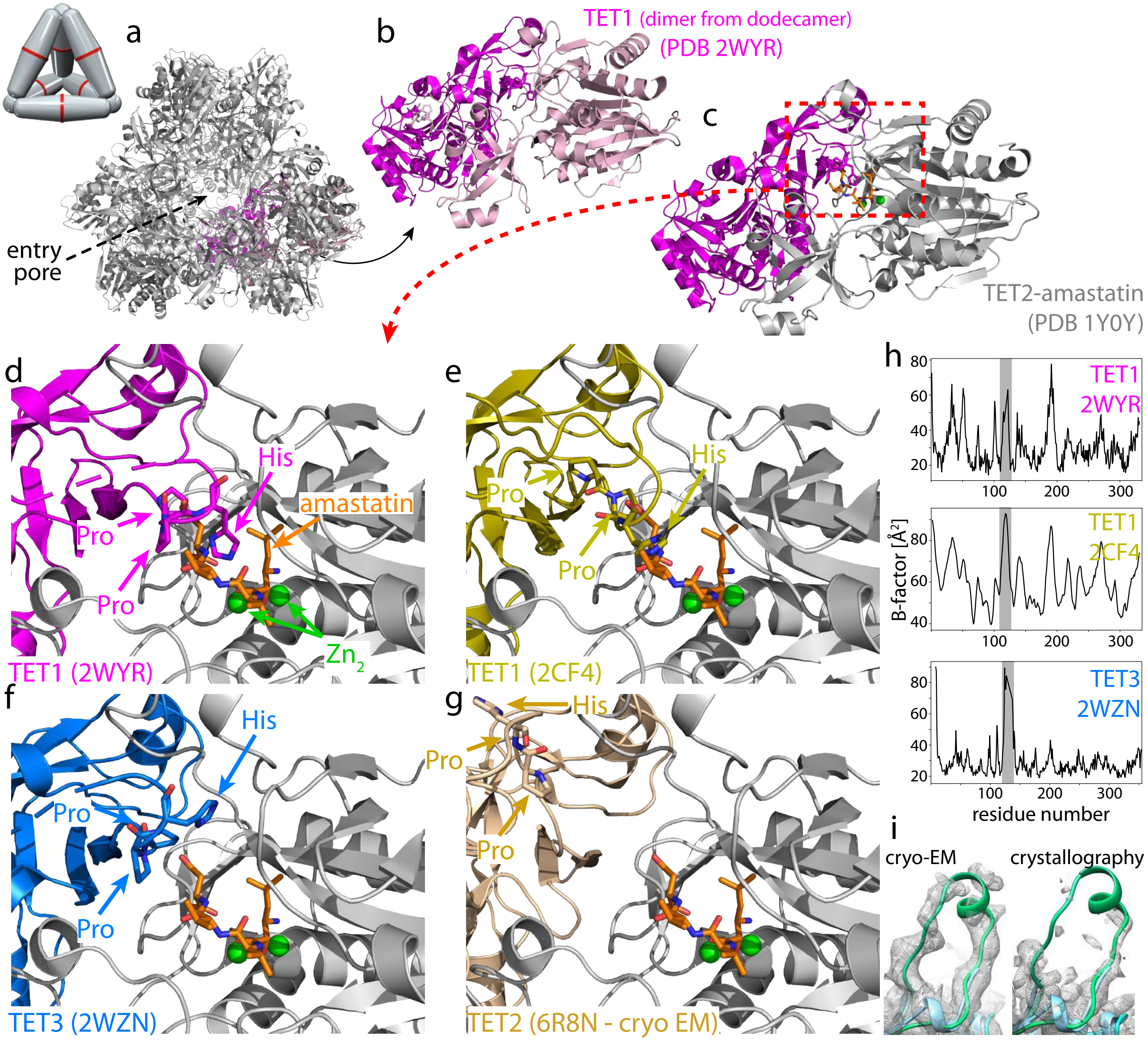


**Fig. S1.** Structures of TET homologs, in which the loop is observed in crystal structures with a focus on the loop. (a) Overview of the dodecamer, highlighting in red/pink two subunits. (b) Zoom onto these two subunits (apo TET1, PDB ID 2WYR). (c) One of the subunits was replaced in silico by the structure of TET2 binding amastatin (PDB ID 1Y0Y, grey). (d)-(g) Zoom onto the active site of TET2 bound to amastatin, and the subunits of TET1, TET3 (crystal structures) or TET2 (cryo-EM/NMR structure from our laboratory). (h) Cα B-factors from crystal structures of apo TET1 and TET3. Note that in TET1 (2wyr), TET2 (1y0r) and TET3 (2wzn) five, thirteen and nine residues of the loop region, respectively, have not been modeled in the crystal structures. (i) Zoom onto the electron density map obtained by cryo-electron microscopy (PDB ID 6F3K) (48) and crystallography (PDB ID 1Y0R) (15). The structure shown as green cartoon has been obtained by a combined NMR/cryo-EM structure calculation (PDB ID 6F3K), whereby the short helix stems from sequence-based TALOS secondary structure prediction. The cyan structure is the crystal structure.

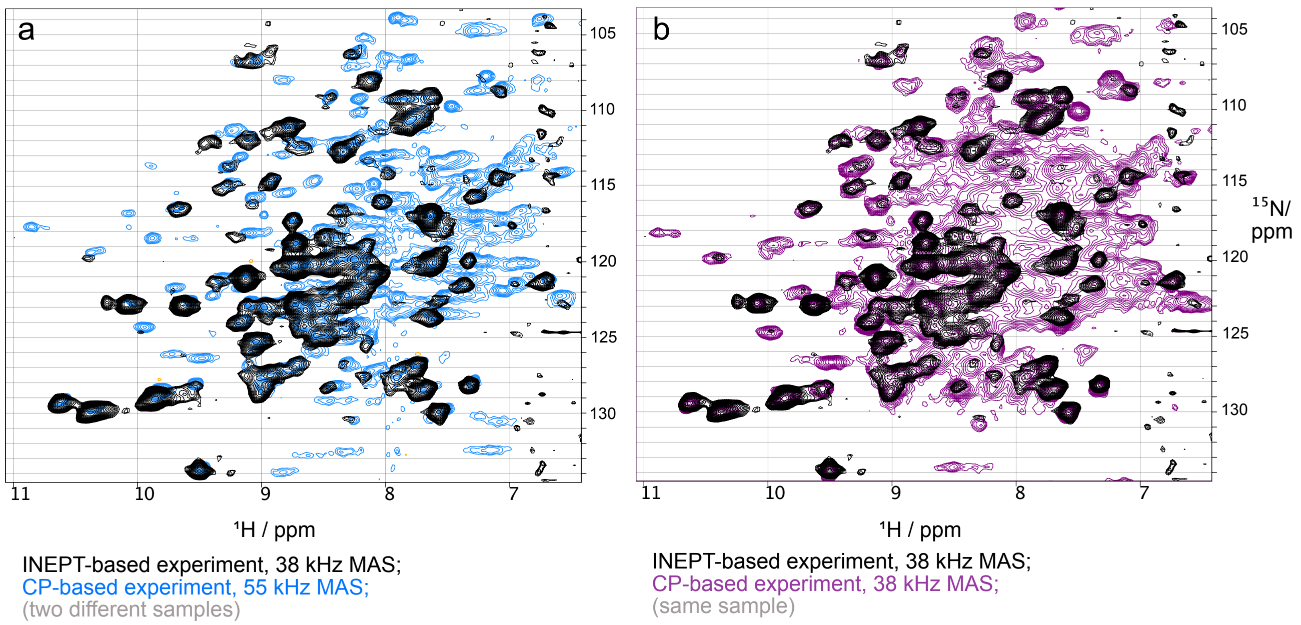


**Fig. S2.** Scalar-coupling based (black) and dipolar-coupling-based (blue, purple) $^{1}$H-$^{15}$N correlation spectra of TET2, prepared with MPD. The scalar-coupling based spectrum is, in principle, able to detect signals for rigid sites as well as for highly flexible sites. In practice, it may fail to do so if the decay of the spin coherences is fast, which may be due to non-averaged dipolar couplings that lead to "coherent decay" (especially at low MAS frequency), or due to the presence of µs-ms dynamics. The dipolar-coupling based experiment is able to detect signals only for sites that are rather rigid, such that the dipolar coupling is not averaged to too small values. It fails for highly flexible sites. Even in deuterated proteins, the decay due to non-averaged dipolar couplings is often fast, such that the scalar-coupling based experiment generally has lower sensitivity (for rather rigid sites) than the dipolar-based one; this trend explains absence of peaks in the scalar-coupling based experiment. The comparison of these experiments may reveal, however, additional peaks in the scalar-coupling based experiment, that may stem from highly flexible sites. This comparison, thus, seeks peaks in the scalar-coupling based experiment that are not present in the dipolar-coupling based one. We recorded the scalar-coupling based experiment at 38 kHz MAS frequency, because at this comparably lower frequency the rigid sites give rise to weaker signals in the scalar-coupling based experiment, increasing the chances to see additional signals. This experiment (black) was compared to dipolar-coupling based (cross-polarization) experiments at two MAS frequencies. The dipolar-coupling based experiment in panel (a) was recorded at 55 kHz MAS (1.3 mm rotor; a different sample than the one of which the scalar-based experiment was obtained), while the one in panel (b) was recorded at 38 kHz, with the same sample used for the scalar-coupling based experiment. We did not detect any additional signals that may be assigned to the loop. A plausible explanation is that µs-ms dynamics make the transfer very inefficient (fast decay). Such µs-ms dynamics is indeed shown in this study for detected methyl sites and the backbone amide of D135. The contour levels in these spectra are on purpose chosen low.

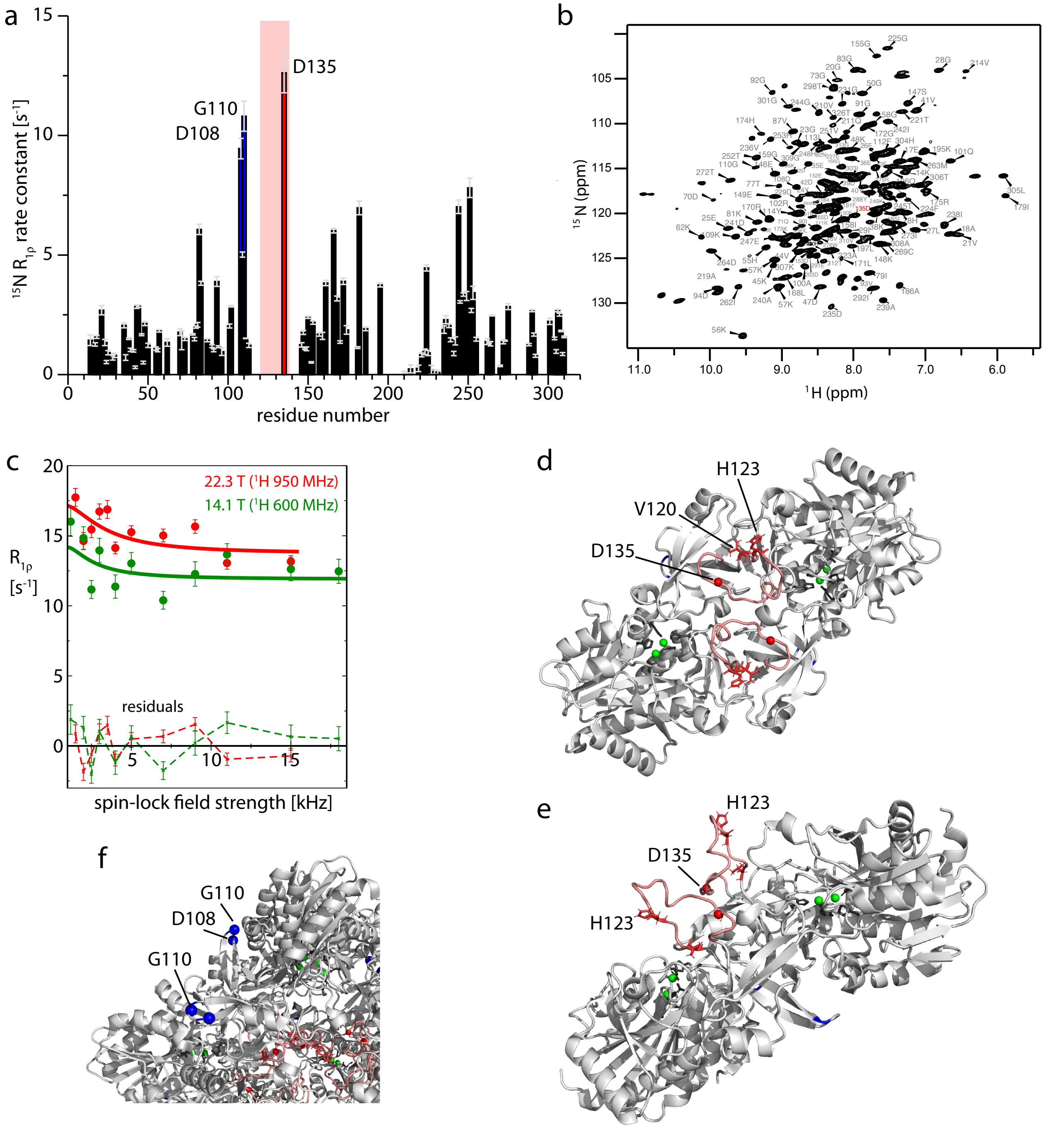


**Fig. S3.** Backbone amide $^{15}$N $R_{1\rho}$ relaxation data, and hint to enhanced loop motion from data of the resolved assigned amide site, D135. (a) $^{15}$N $R_{1\rho}$ rate constants obtained with a perdeuterated, 100% back-exchanged TET2 sample spinning at 39 kHz MAS frequency, at a spin-lock field strength of 15 kHz (14.1 T magnetic field strength). Highlighted in red is the data of residue D135. The loop region (120-138) is highlighted in red. (b) Two-dimensional $^{1}$H-$^{15}$N correlation spectrum (CP-based) at 55 kHz MAS frequency, as reported in (48). The peak assigned to D135 is highlighted with a red label. (c) $^{15}$N $R_{1\rho}$ Bloch-McConnell relaxation dispersion profiles of D135. Non-flat profiles are indicative of structural fluctuations that involve a variation of the chemical shift. The data have been obtained at two different static magnetic field strengths (14.1 and 22.3 T) as indicated. The solid lines are fits of both data sets to a 2-site exchange model that implies a kinetic rate constant and the product of the populations of the two states and their squared chemical-shift difference. The precision of such a fit to a single residue is generally poor, and the two-site exchange model certainly not appropriate here; these lines are used here only to highlight that there is µs dynamics involving D135. (d, e) Position of D135 within the loop (red sphere), as well as the important residue H123 and the methyl probe V120. The zinc atoms in the active site (green) and the chelating residues (black) are shown on this dimer, out of the dodecameric assembly. (f) Zoom onto the two other residues that show high $^{15}$N $R_{1\rho}$ rate constants, D108 and G110; these residues are in surface exposed loops, and are unrelated to the catalytic chamber.

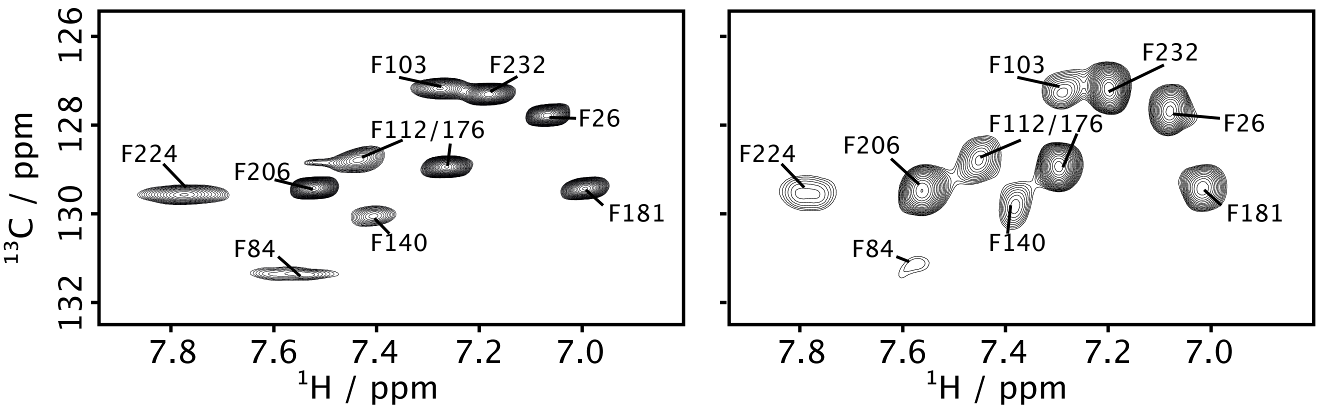


**Fig. S4.** $^{1}$H-$^{13}$C MAS NMR correlation spectrum of a specifically labeled H123F TET2 sample (right) and a WT TET2 sample (left), akin to spectra reported earlier (50). The labeling scheme comprised uniform deuteration and $^{15}$N labeling as well as $CHD_2$-groups at Leu and Val (stereospecific labeling of the pro-R site) and the $^{1}$H-$^{13}$C pair at the para-CH position of Phe. The spectrum of the H123F variant shows ten peaks, corresponding to the native phenylalanines, but no additional peak that may be assigned to the non-native F123. We ascribe the broadening of the signal to µs dynamics of the loop and its resulting enhanced relaxation. This data is in agreement with the backbone H-N directed spectra, in which most sites of the loop are not visible, and the methyl data, which show strongly enhanced relaxation of V120 and I139. Note that there are small shifts in H123F for signals belonging to F84, F140 (in immediate vicinity to each other) and a minor shift of F103. We ascribe those to a small change in temperature. The difference in line width is explained by different acquisition parameters, which were optimized in the H123F case towards higher sensitivity (rather than resolution), and is inconsequential for this comparison.

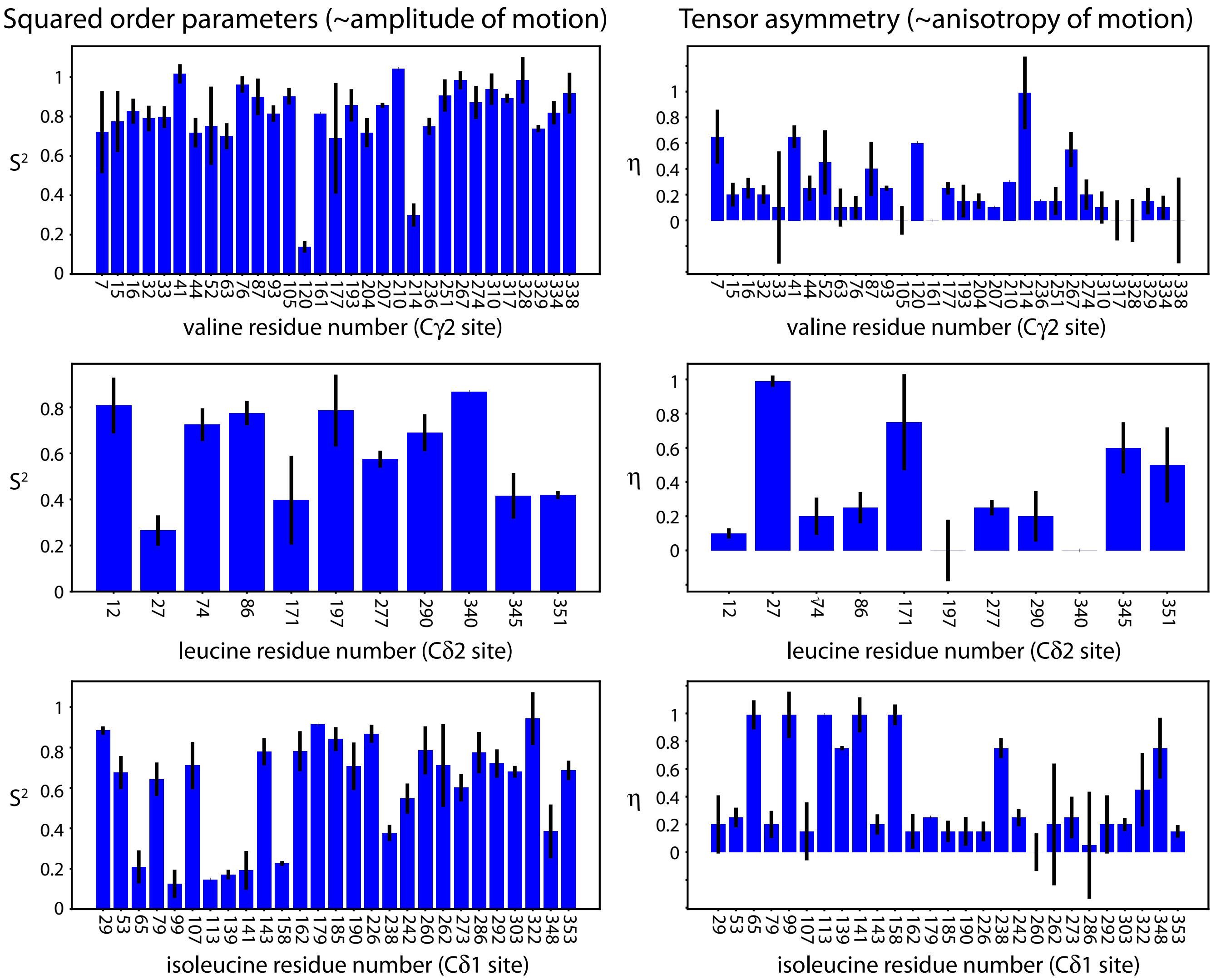


**Fig. S5.** Results of REDOR fits of Ile-$\delta$1, Leu-$\delta$2, Val-$\gamma$2 $CHD_2$ methyl sites in TET2. Left: squared dipolar order parameters obtained as the square of the ratio of the measured $^{1}$H-$^{13}$C dipolar-coupling anisotropy over the rigid-limit value 14529 Hz (taking into account that the methyl rotation reduces it by three-fold from the value obtained for a 1.115 Åbond length). Right: dipolar-coupling anisotropies.

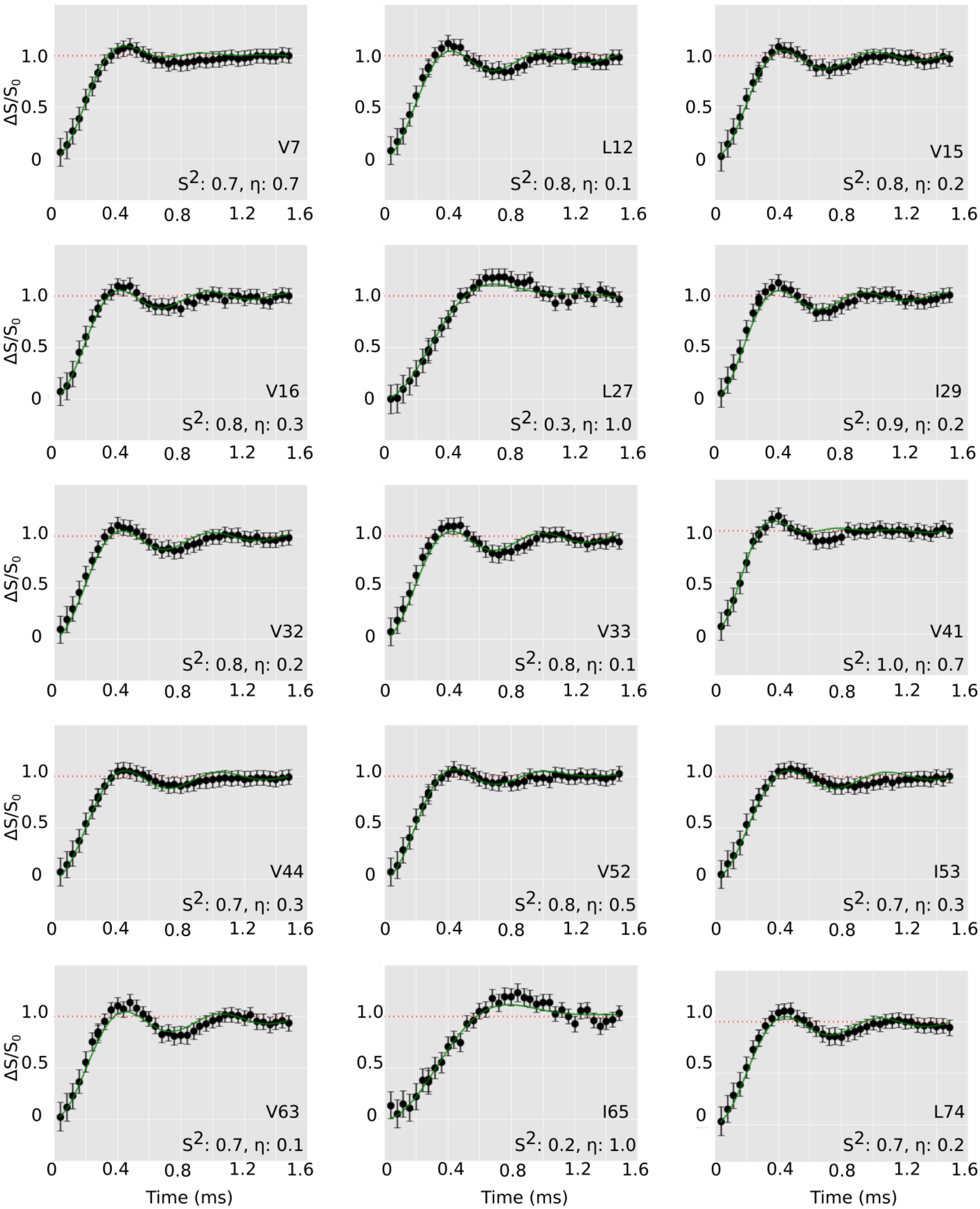


**Fig. S6.** $^{1}H$-$^{13}C$ REDOR curves for $CHD_2$ methyl groups in apo TET2 at 55.555 kHz MAS frequency. Solid lines show the best-fit REDOR curves, in which the anisotropy of the dipolar-coupling tensor (from which the order parameter is calculated) and the tensor asymmetry are the only fit parameters.

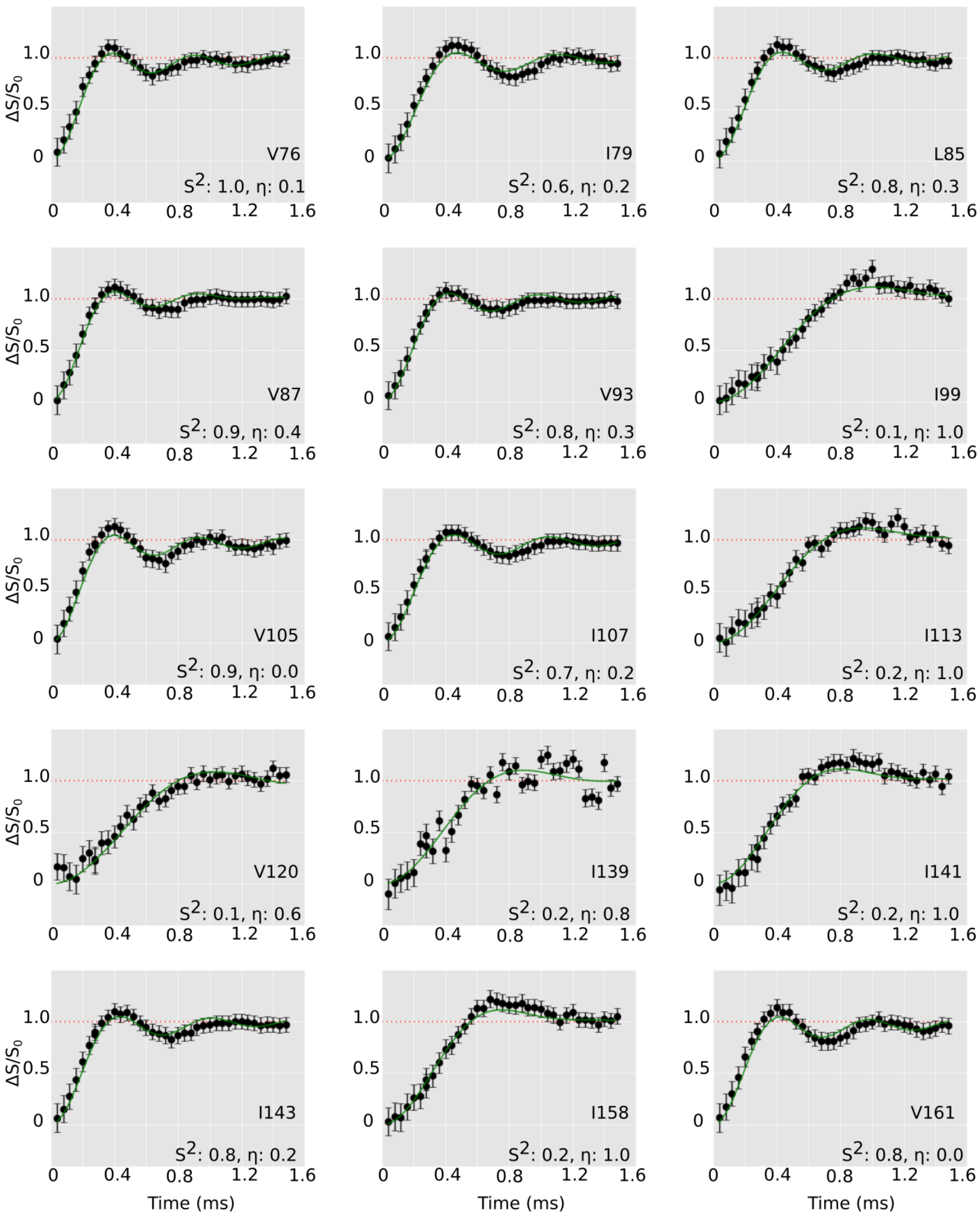


**Fig. S6.** (continued) $^{1}H$-$^{13}C$ REDOR curves for $CHD_2$ methyl groups in apo TET2.

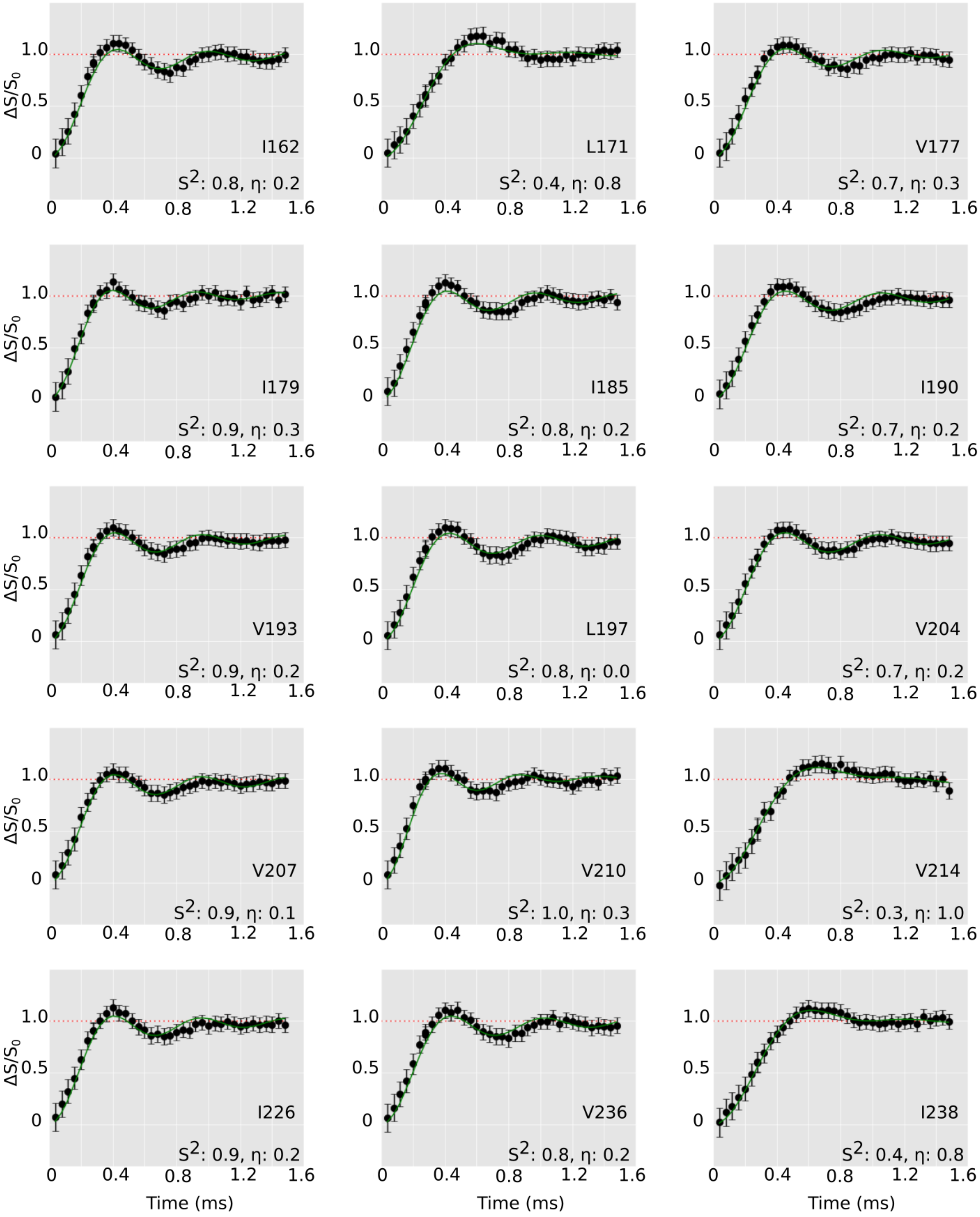


**Fig. S6.** (continued) $^{1}$H-$^{13}$C REDOR curves for $CHD_2$ methyl groups in apo TET2.

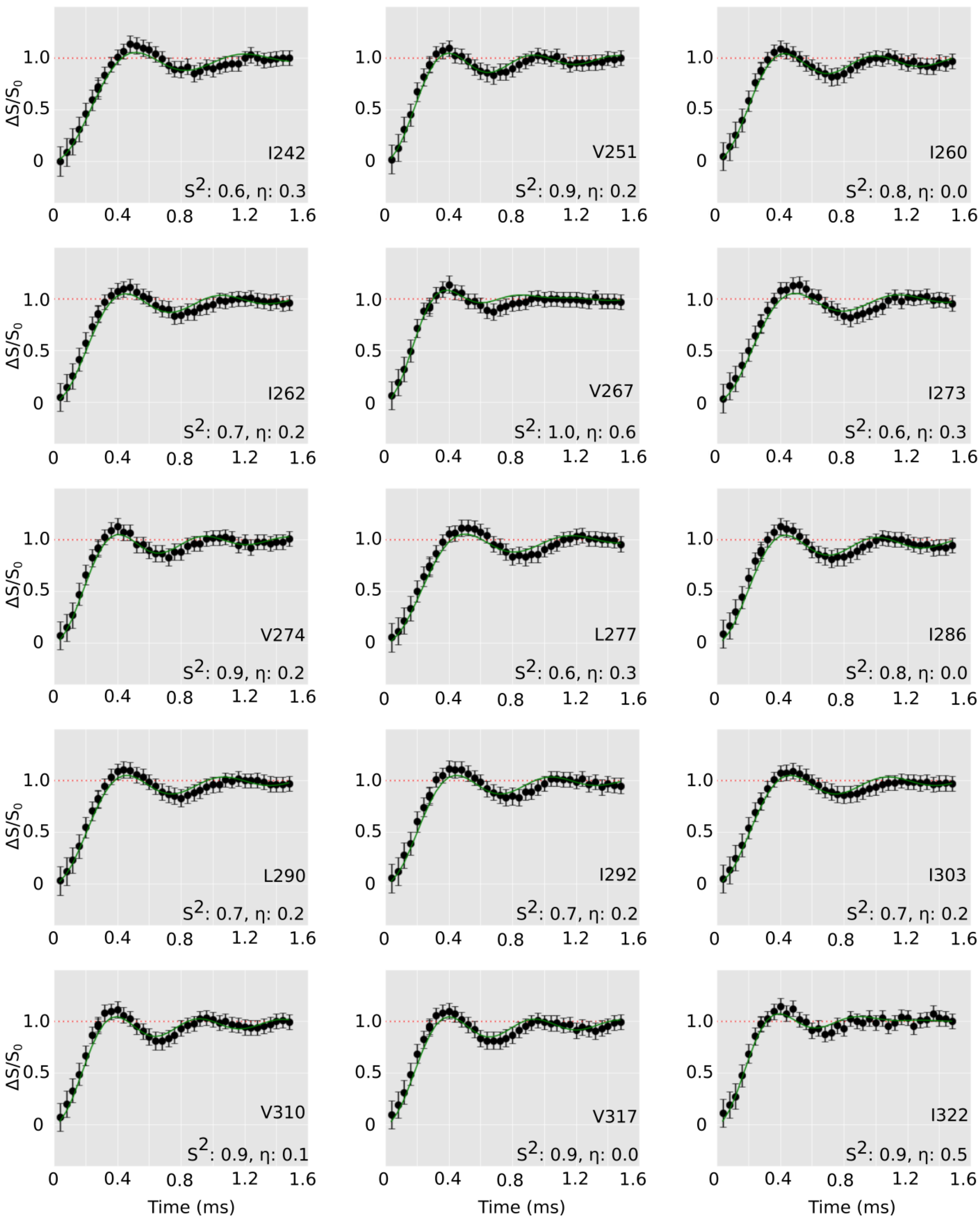


**Fig. S6.** (continued) $^{1}H$-$^{13}C$ REDOR curves for $CHD_2$ methyl groups in apo TET2.

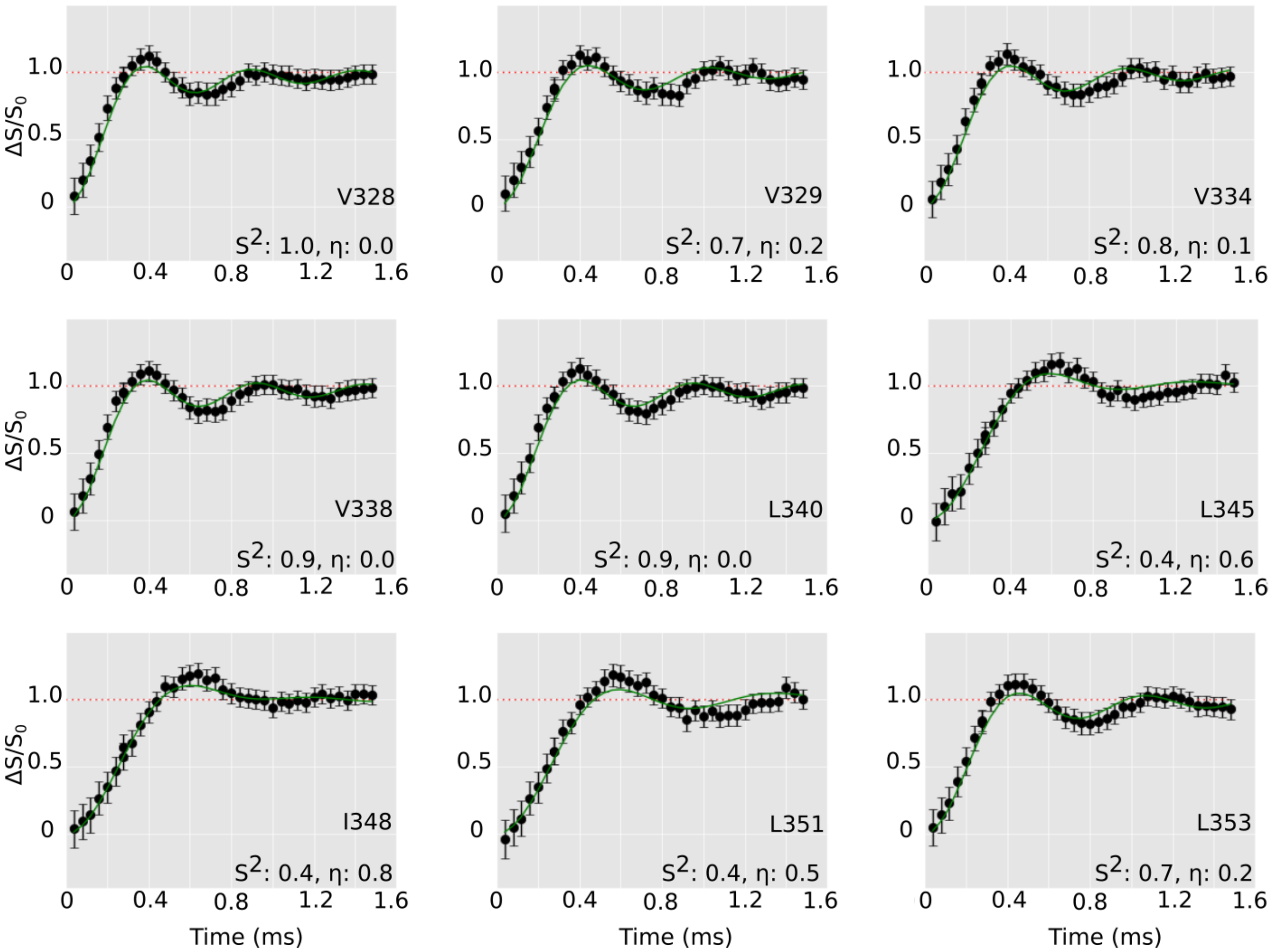


**Fig. S6.** (continued) $^{1}H$-$^{13}C$ REDOR curves for $CHD_2$ methyl groups in apo TET2.

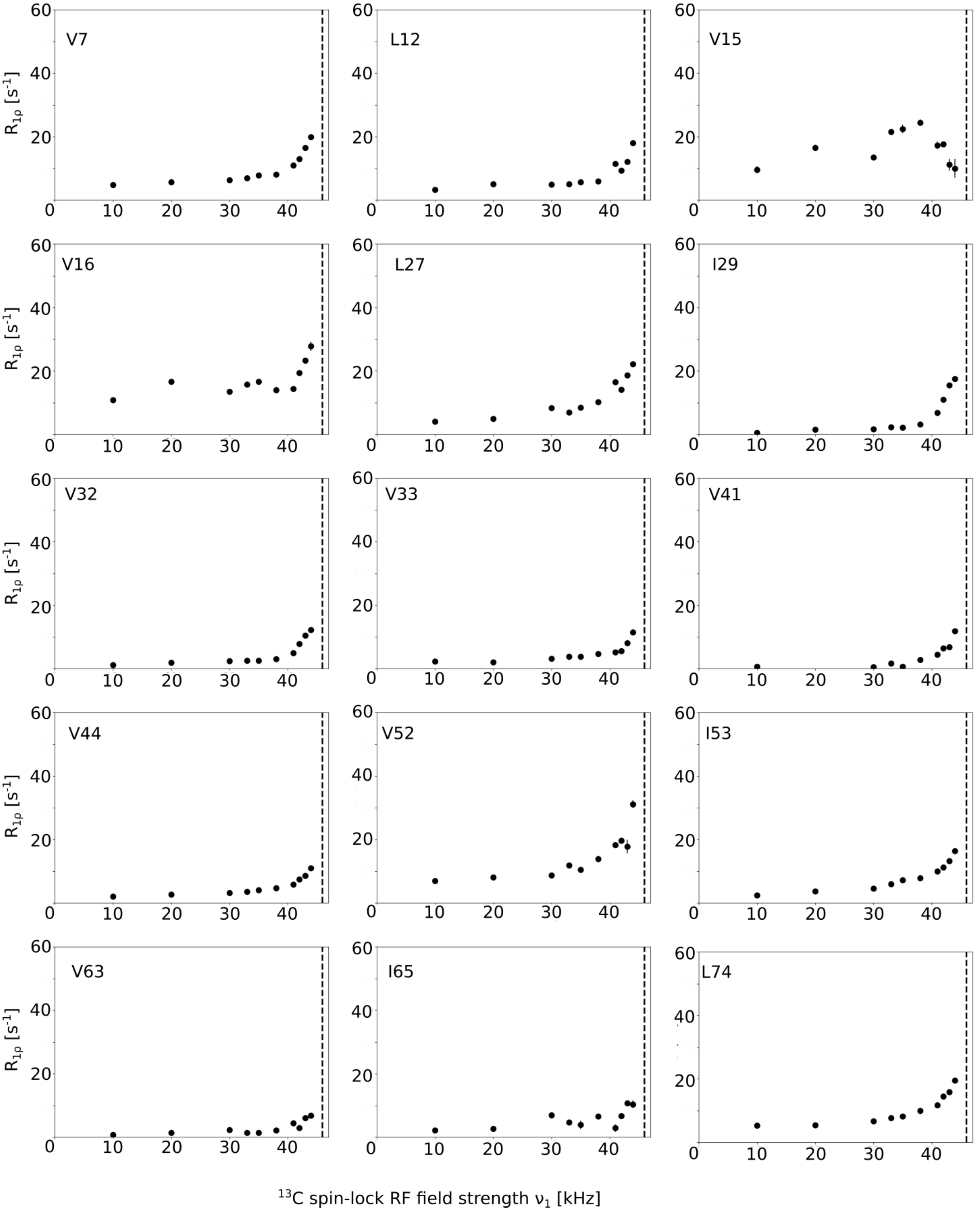


**Fig. S7.** $^{13}$C $R_{1\rho}$ NERRD curves for $CHD_2$ methyl groups in apo TET2. Each data point corresponds to the rate constants obtained from an exponential fit of peak intensities as a function of the spin-lock duration (see Methods). The vertical line indicates the RF field strength at which the nutation frequency matches the MAS frequency (i.e. the n=1 rotary-resonance condition).

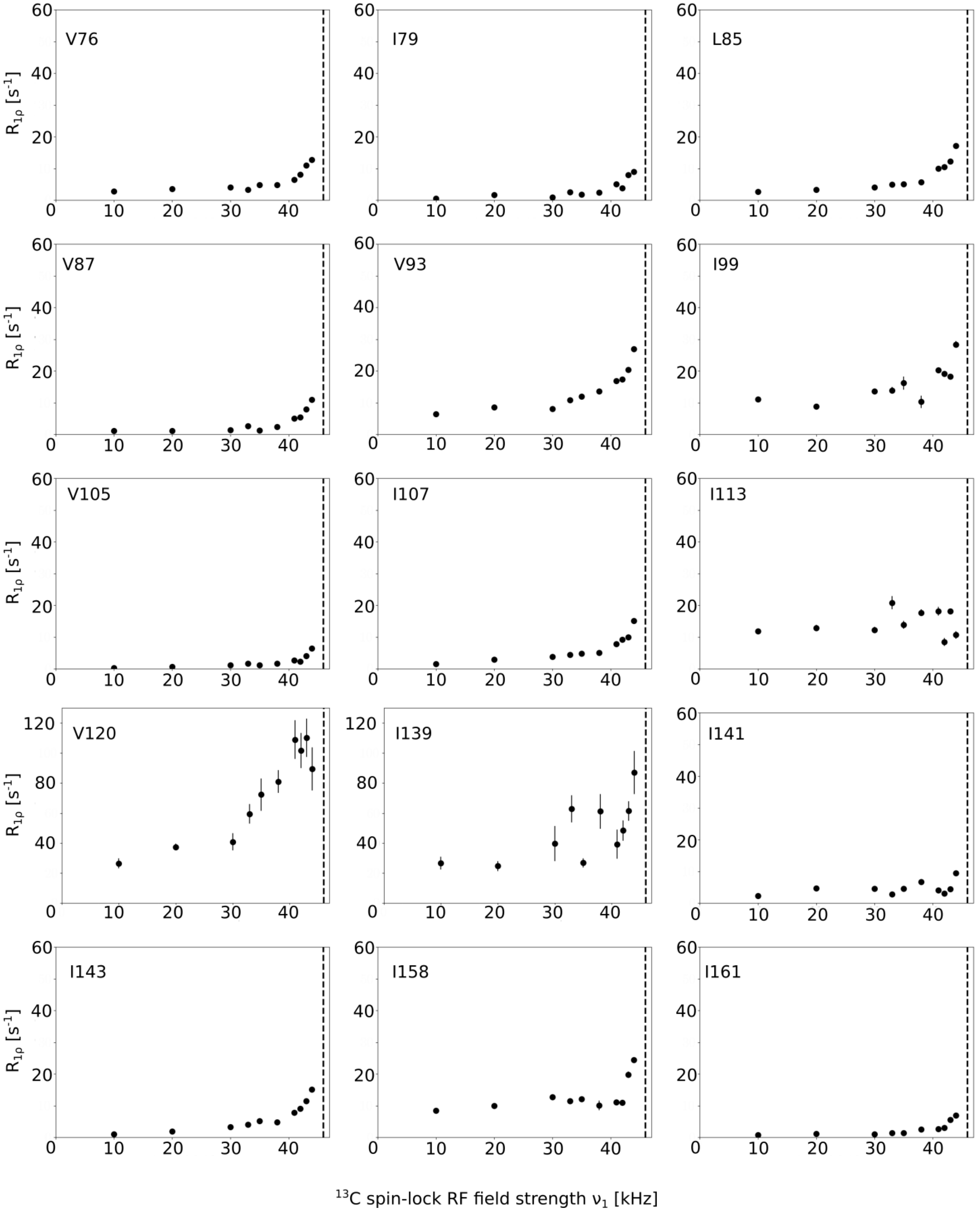


**Fig. S7.** (continued) $^{13}$C $R_{1\rho}$ NERRD curves for $CHD_2$ methyl groups in apo TET2.

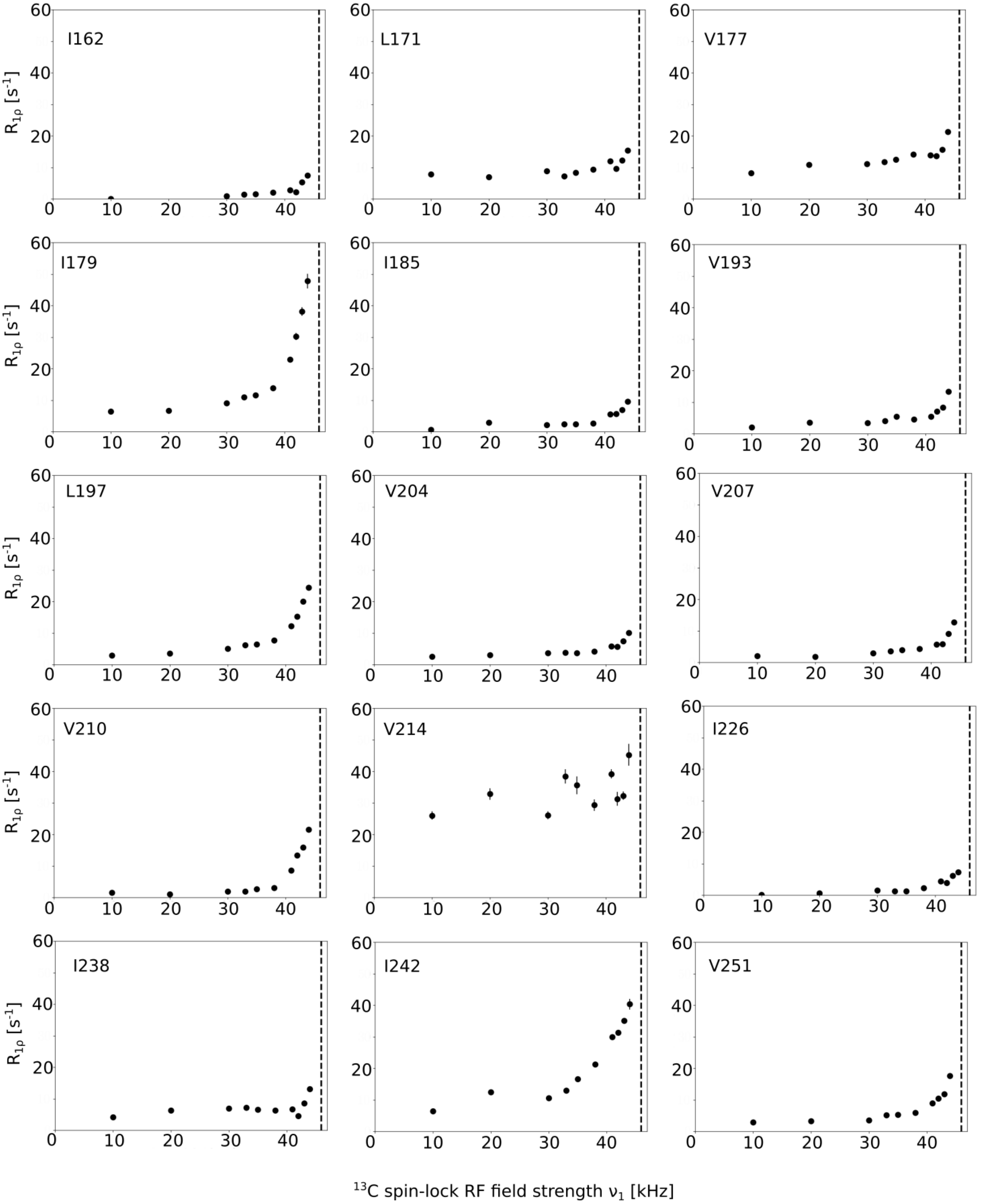


**Fig. S7.** (continued) $^{13}$C $R_{1\rho}$ NERRD curves for $CHD_2$ methyl groups in apo TET2.

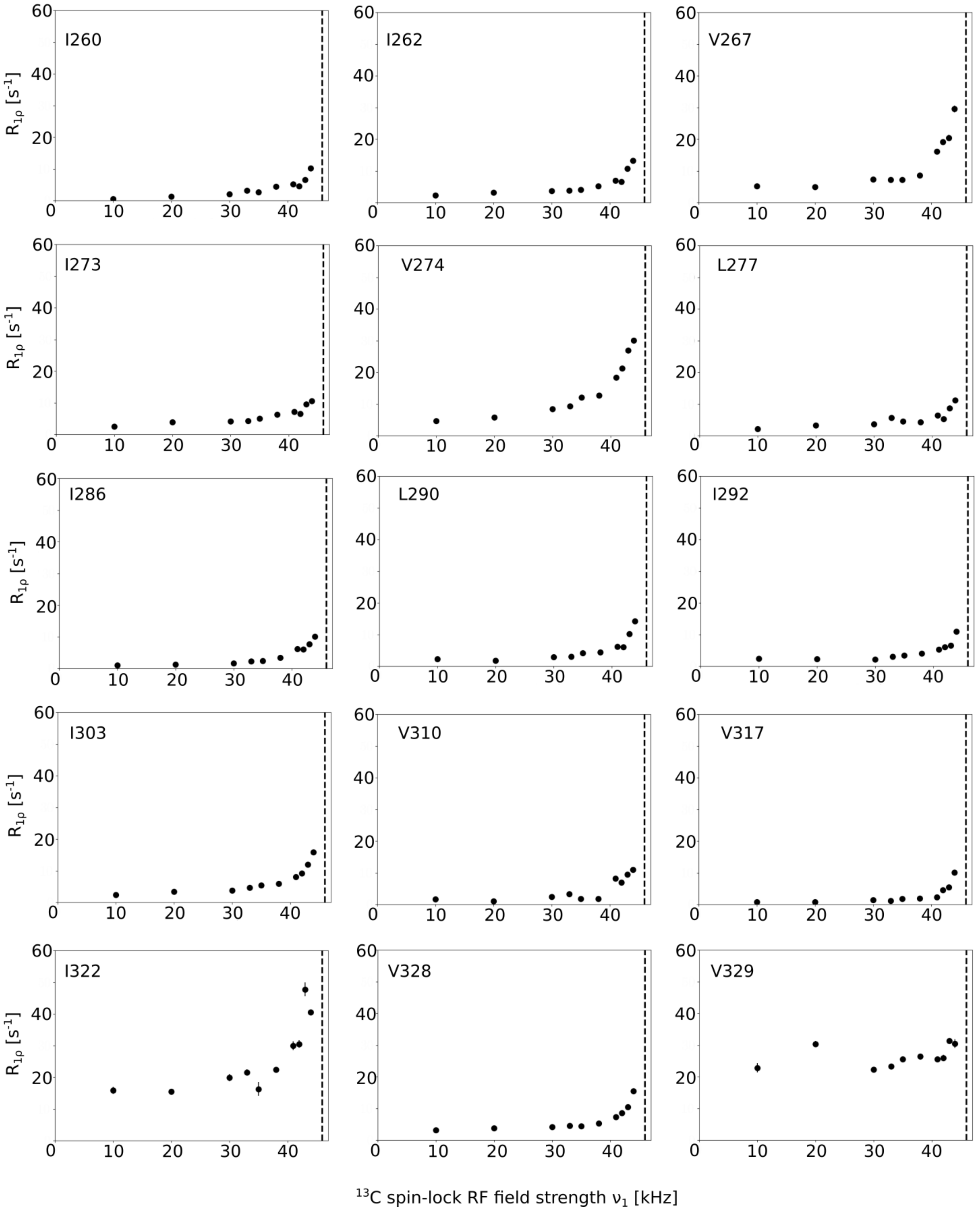


**Fig. S7.** (continued) $^{13}$C $R_{1\rho}$ NERRD curves for $CHD_2$ methyl groups in apo TET2.

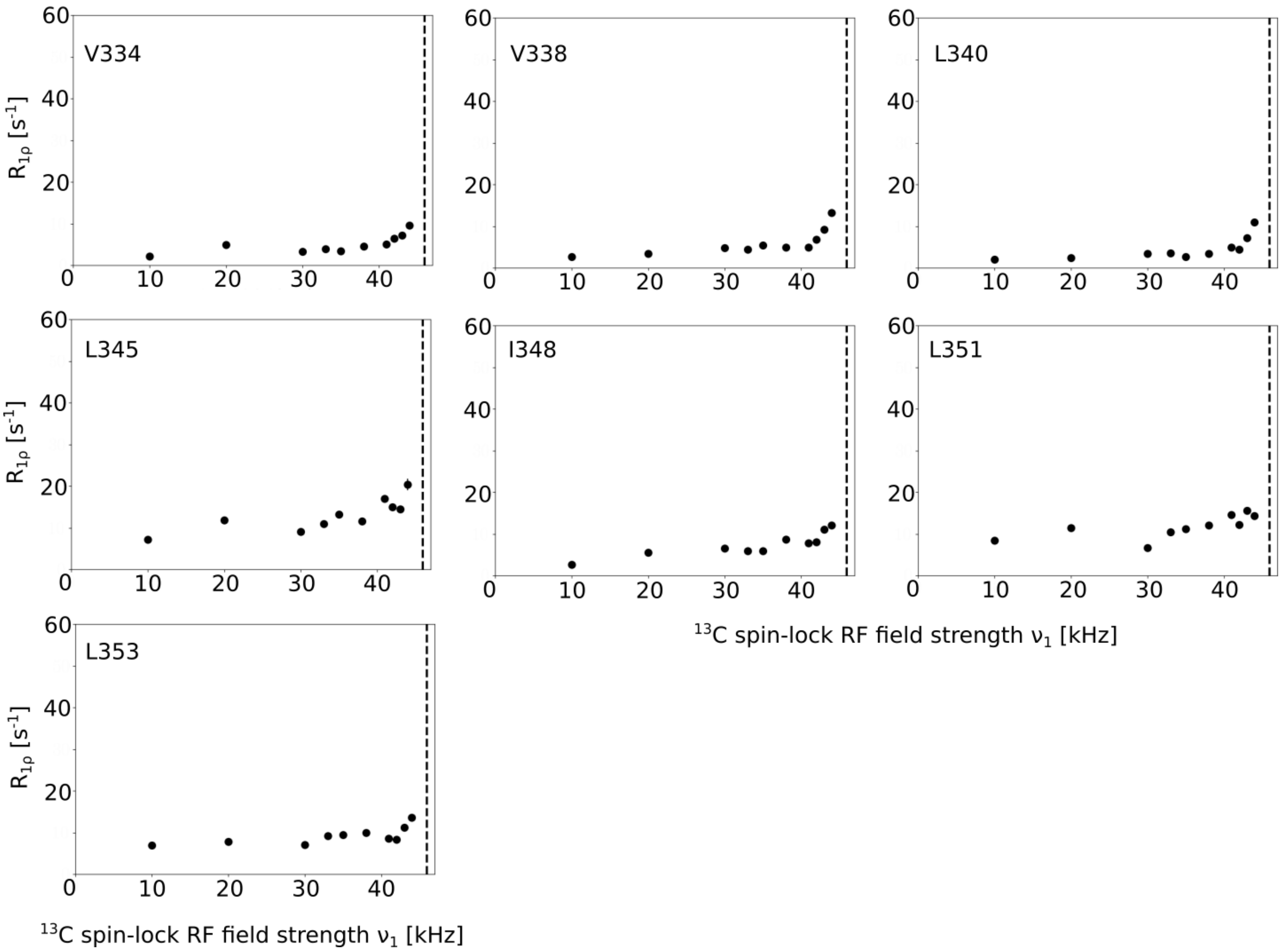


**Fig. S7.** (continued) $^{13}$C $R_{1\rho}$ NERRD curves for $CHD_2$ methyl groups in apo TET2.

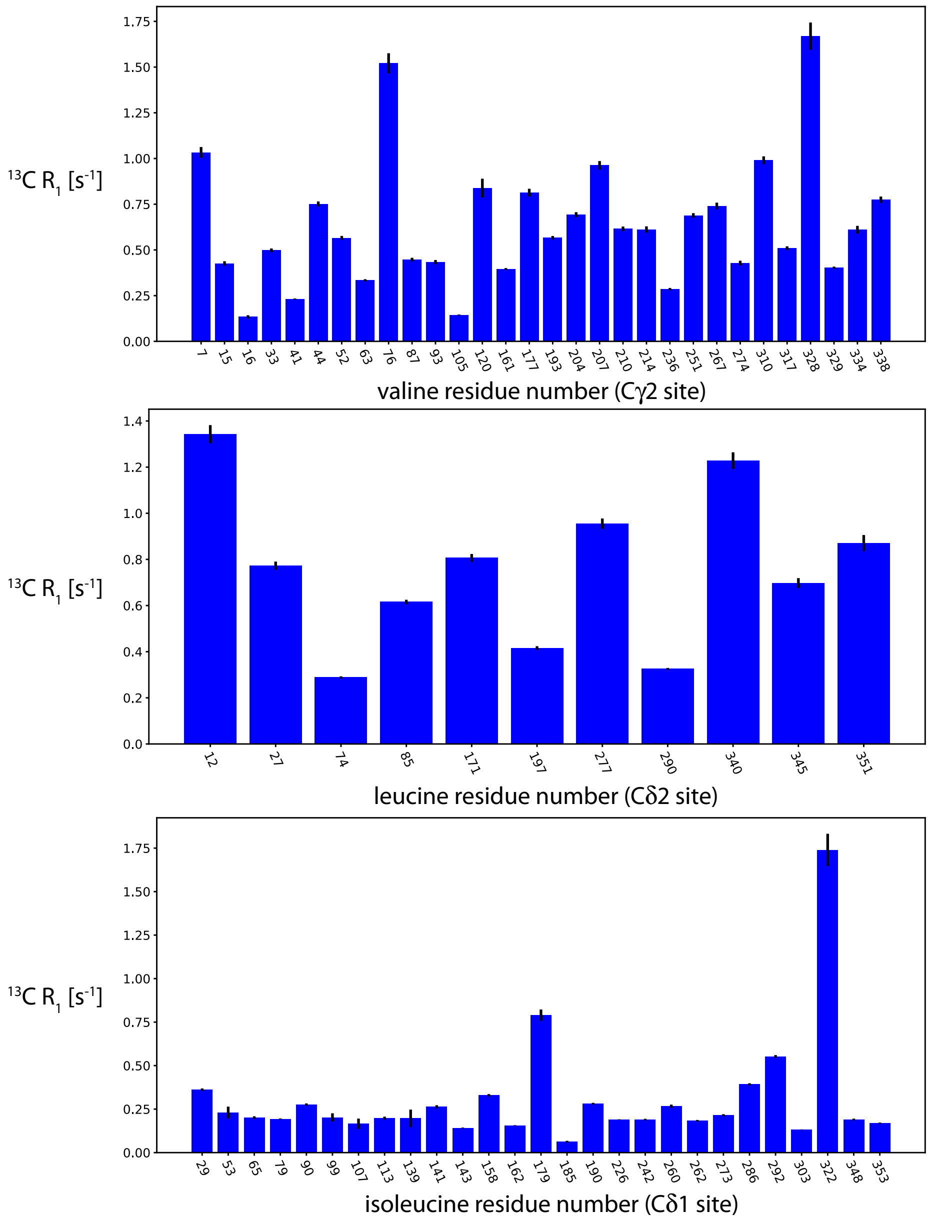


**Fig. S8.** $^{13}$C longitudinal relaxation rate constants, $R_1$, measured at a $B_0$ field strength of 22.3 T. The $R_1$ rate constant is particularly sensitive to nanosecond motion (52, 58). The methyl sites of V120 and I139 do not have particularly high values, suggesting that the loop motion is not on the nanosecond time scale, but - as revealed by the $^{13}$C $R_{1\rho}$ data - on µs time scales.

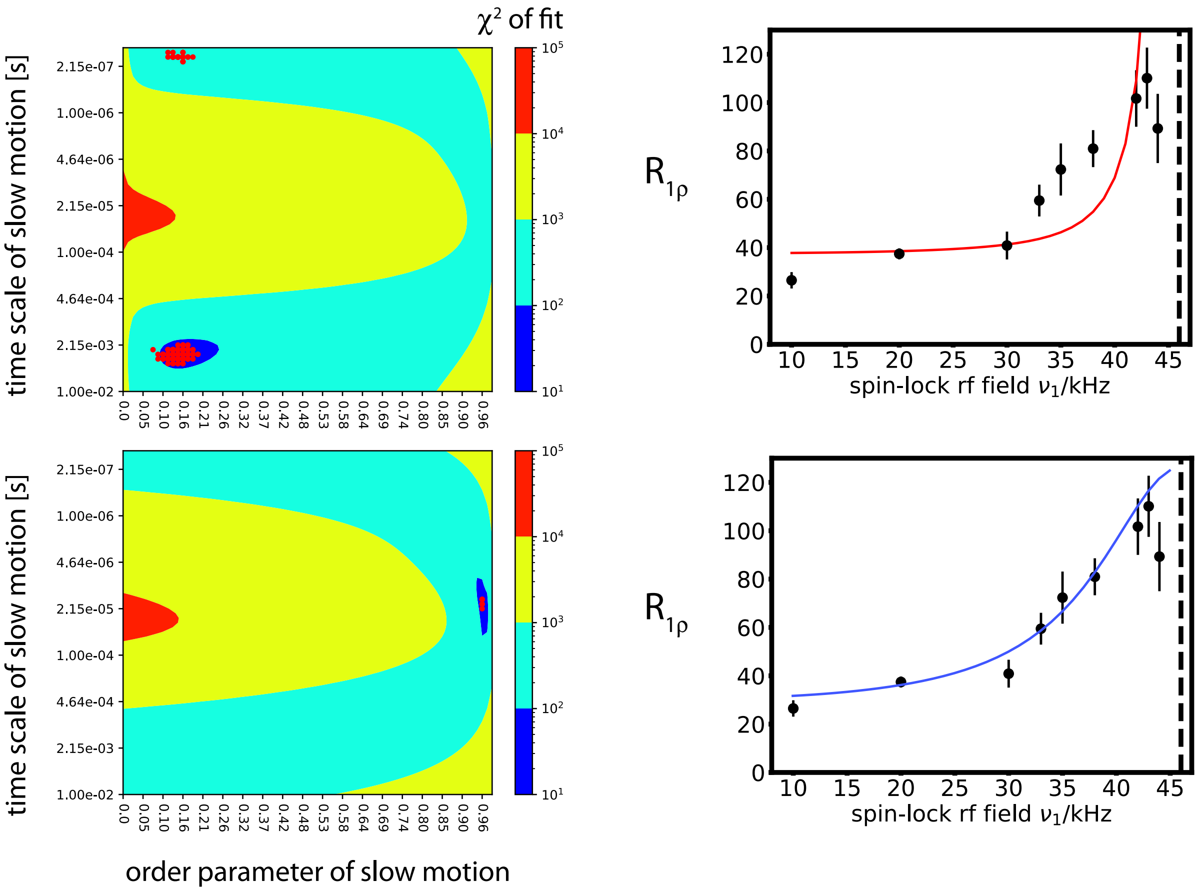


**Fig. S9.** Outcome of the fit of a simple motional model of the V120 relaxation and dipolar coupling data. The model comprises fast methyl rotation with a squared order parameter of 1/9 and a "fast" time scale correlation time, as well as a slow motion with a variable order parameter and a variable correlation time. Two different fit scenarios are shown, whereby either the "slow" order parameter was fixed to match the REDOR-derived value (red curve and upper chi-square plot), or the REDOR data were not used, and only the $^{13}$C $R_{1\rho}$ values, shown in the right graph, and the $^{13}$C $R_1$ value were used. The corresponding time scales of the slow motion in these two scenarios range from ca. 10 µs to 1 ms. The fact that the better fitting curve (blue) leads to an order parameter which is not in agreement with the REDOR order parameter strongly suggests that the motional model is too simplistic. In the context of this study, the exact time scale of motion is not important and we refrained from more complex models, rather limiting the interpretation to saying that the time scale is in the ca. 10-1000 µs range.

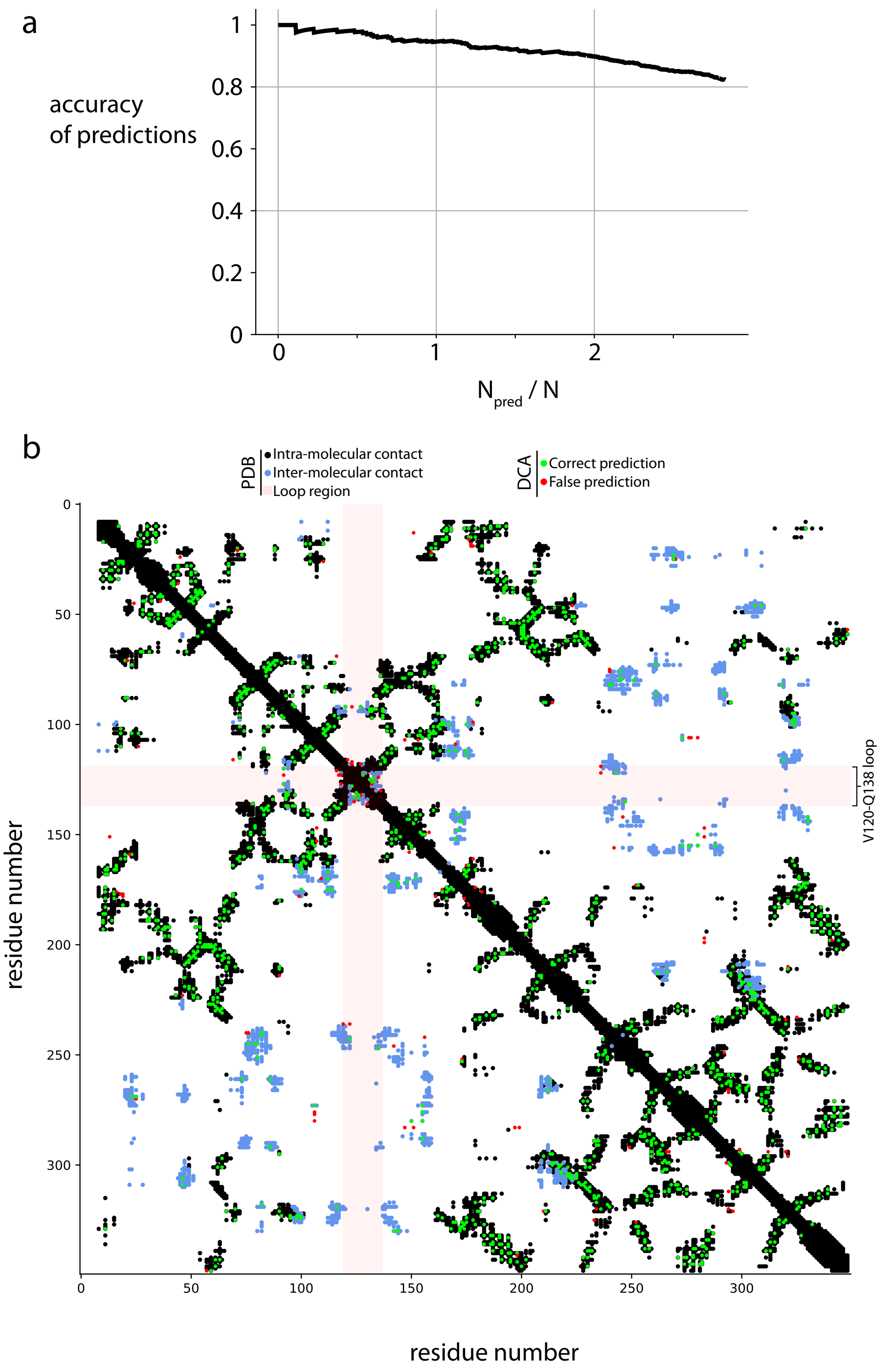


**Fig. S10.** Direct Coupling Analysis (DCA) of TET2. (a) DCA prediction accuracy curve, determined from the structure (PDB: 6R8N). Structural contacts were defined by inter-atomic distances between heavy-atoms below 8.5 Å. The accuracy is shown as a function of the number of top predictions, normalized by the total number of positions in the sequence, N=353. DCA prediction show excellent prediction accuracy over a large range of predictions. Notably, considering the top 2N=706 highest ranked DCA predictions results in a prediction accuracy of 88%. Ignoring the false positives rising from predictions falling in regions where the PDB structure is not defined, the accuracy rises above 90%. (b) Residue-residue contact plot. Shown in red/green are the top 2N (706) highest ranked inter-residue contacts predicted by DCA (red: false positives, green: true positives). Predictions are overlaid on and compared to the structural contact map obtained from an experimentally determined model (PDB: 6R8N). Intra-molecular structural contacts are depicted in black, inter-molecular structural contacts in blue. The residue-positions numbering relates to the reference sequence of the *Pyrococcus horikoshii* TET2 Peptidase (UniprotID O59196). Structural inter-residue contacts are defined if the inter-atomic distance for any pairs of atoms in two residues is below 8.5Å.

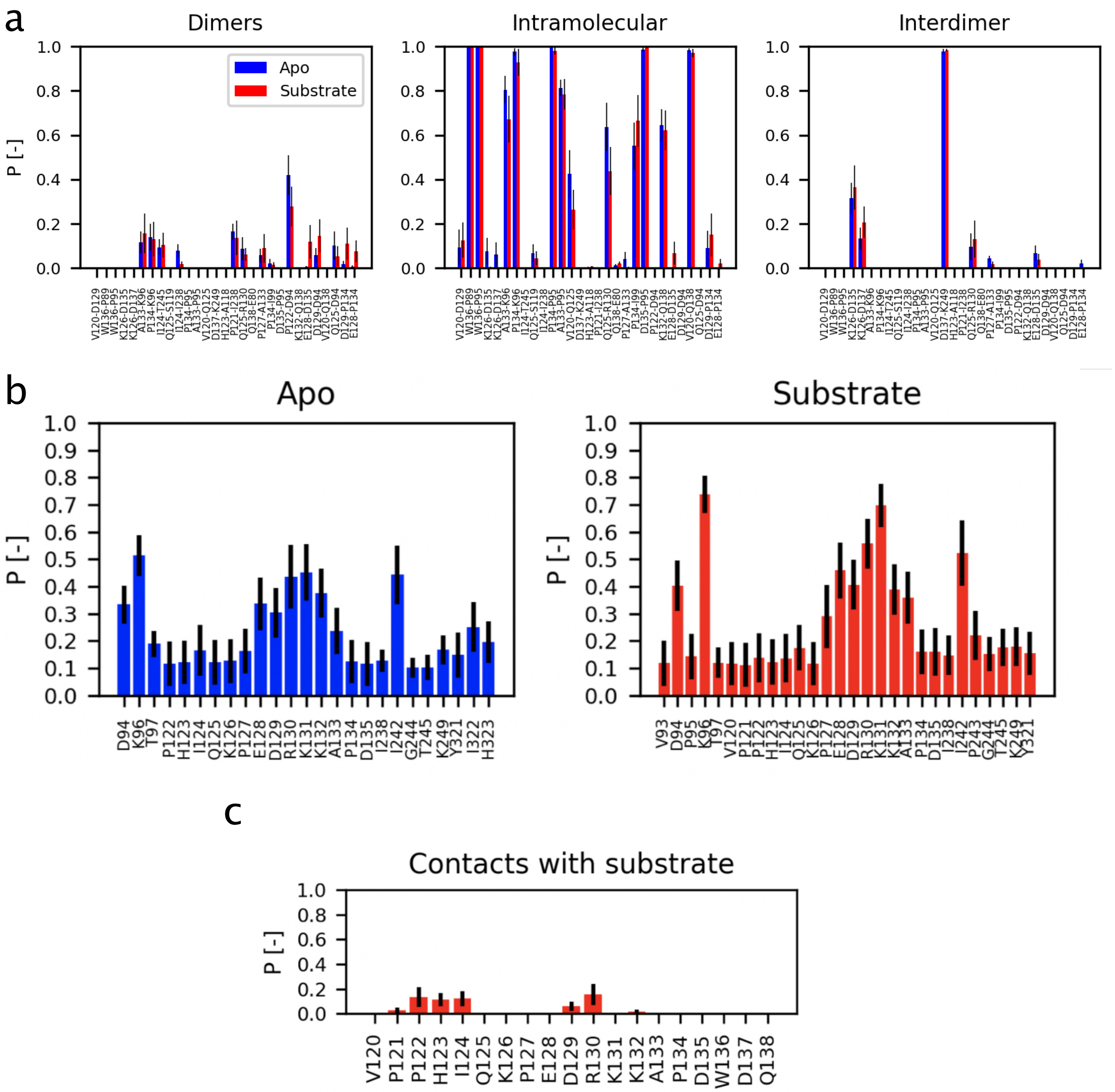


**Fig. S11.** Contacts observed in the 1 µs molecular dynamics simulation of the full dodecameric assembly, plotted as probability, i.e. fraction of time during which the contacts are present. (a) Observation of residue-residue contacts either in between two adjacent subunits (i.e. within one of the six dimers that form the TET2 assembly; left), or within the same subunit (intramolecular; center), or from one loop to a loop belonging to a subunit outside that dimer. (b) Identification of the residues with which the loop is in contact, either in the apo state (left) or the substrate-bound state (right). (c) Contacts formed by residues of the loop with a substrate (Leu-Leu-Val-Ala) in the active site. In all cases, a contact is defined as having the minimum distance between two heavy atoms of the concerned residues below 5 Å.

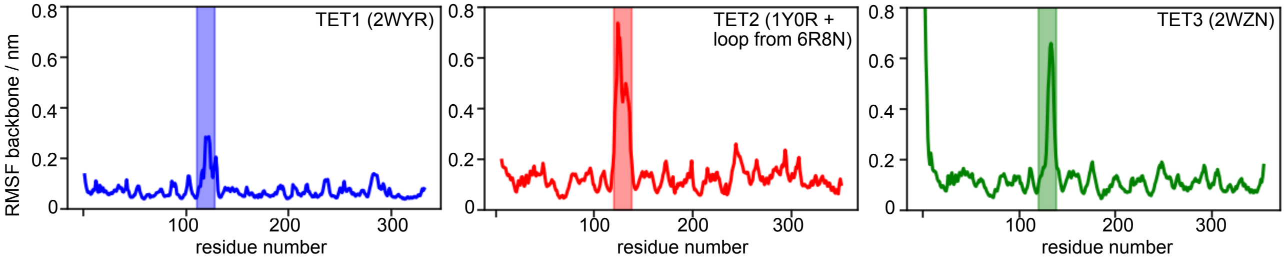


**Fig. S12.** The loop is dynamic in all three archaeal TET assemblies, as probed by 1 µs long MD simulations of TET1, TET2 and TET3. (In contrast to Figure S11, only the dimers were simulated here, in order to sample sufficient time scales within reasonable times.) In these simulations backbone is constrained since dimers are not supposed to be thermodynamically stable, the loop and neighbouring regions ($\pm$ 10 residues) are free to move. In all cases, the loop is dynamic, although somewhat less in the simulation of TET1. This observation may be due to sampling limitations and diverse starting conformations: in the TET1 X-ray structure the loop is much closer to the active site.

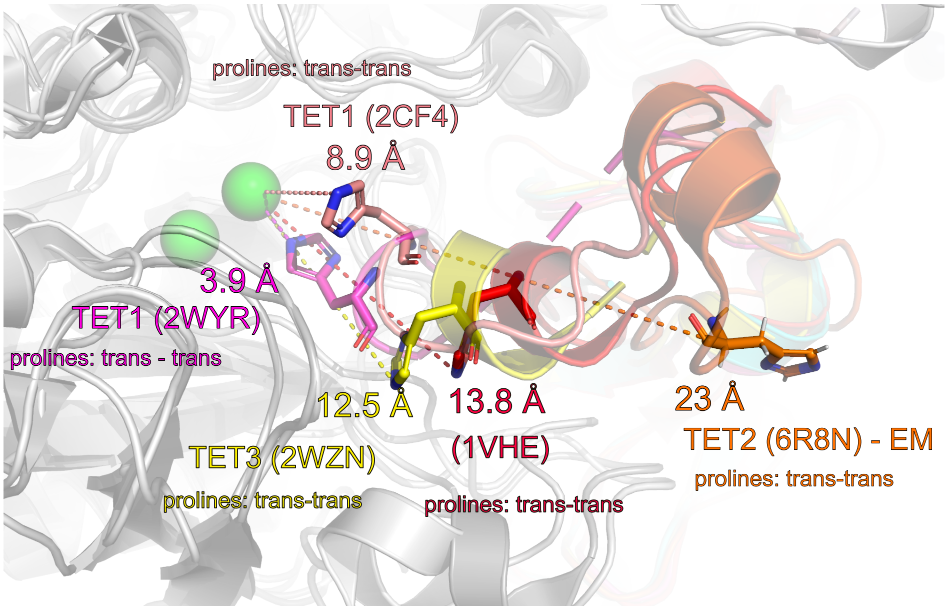


**Fig. S13.** Close-up of the distance between the conserved histidine in the loop of one subunit and the active site of the adjacent subunit. In all cases, the subunit of which the loop is considered is shown in color, while the adjacent subunit is shown in grey, with the active-site zinc ions in green. The identity of the proteins and the PDB access codes are written in the figure. The structure with PDB code 1VHE is a homolog from *B. subtilis.* Further homologs, which are not displayed here for reasons of readability of the figure show similarly long distances between the His and the zinc site. In PfTET4 (PDB 4x8i) the distance is ca 11 Å , and in a bacterial homolog (PBB 4wwv) it is ca. 12 Å. We have also analyzed the Pro-Pro sequence that precedes the His in all these proteins, in particular their cis/trans conformation. As indicated, all prolines are in trans conformation.

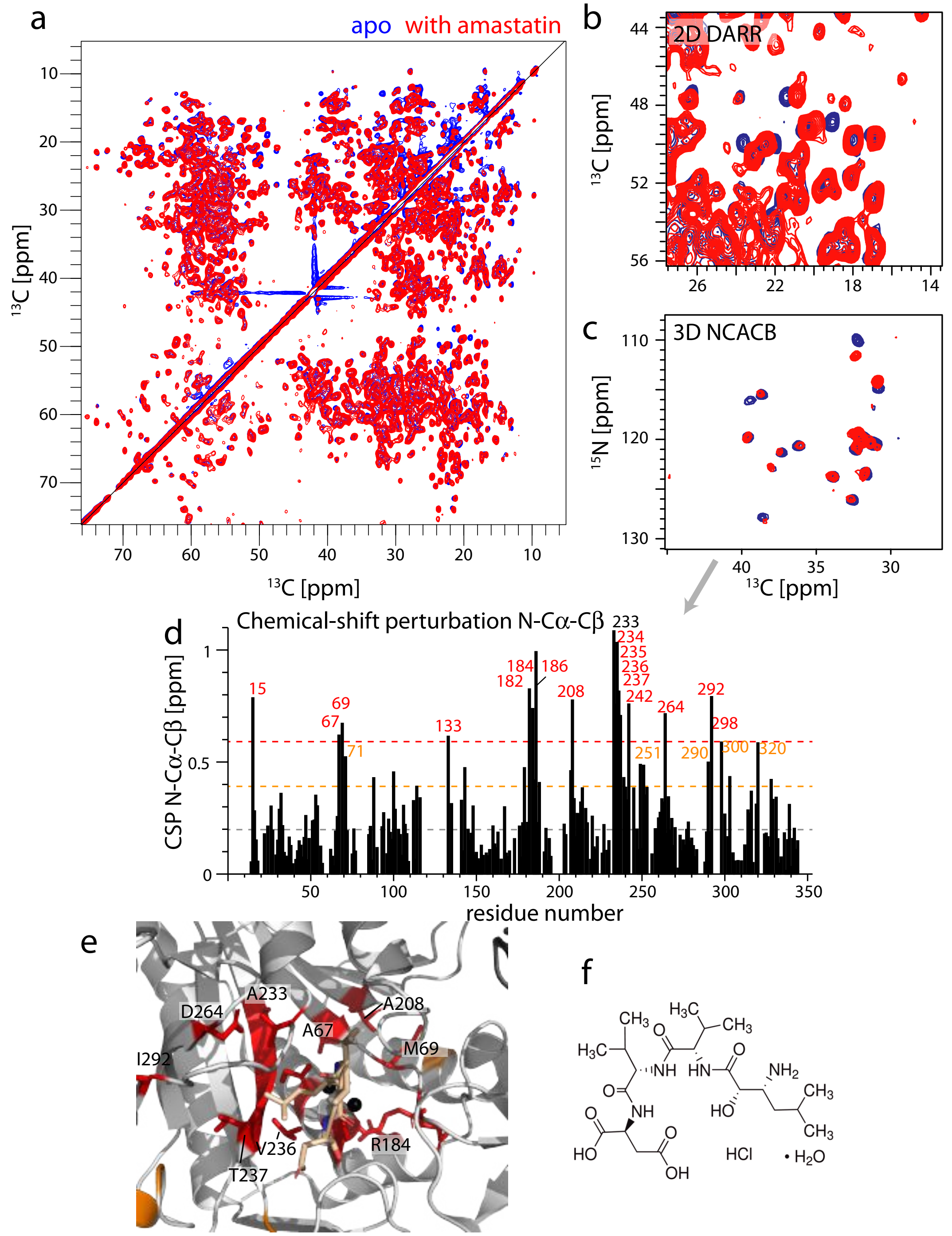


**Fig. S14.** Amastatin binding in TET2 observed by $^{13}$C-detected MAS NMR. (a) 2D $^{13}$C-$^{13}$C DARR correlation spectra of u-[$^{13}$C,$^{15}$N]-labeled TET2 precipitated by addition of 2-methyl-2,4-pentanediol (MPD) to a solution without amastatin (blue) and after having incubated the protein with an excess of amastatin (red) at 15 kHz MAS frequency. (b) Zoom of the spectrum in (a). (c) 2D projection of the 3D NCACB spectrum, projected along the Cα dimension from 63 to 68 ppm. (d) Quantitative analysis of the chemical-shift perturbations (CSPs) of the N, Cα and Cβ frequencies. The number of the residues with the most important CSPs are indicated and plotted onto the structure of TET2 in panel (e). (f) Chemical structure of amastatin.

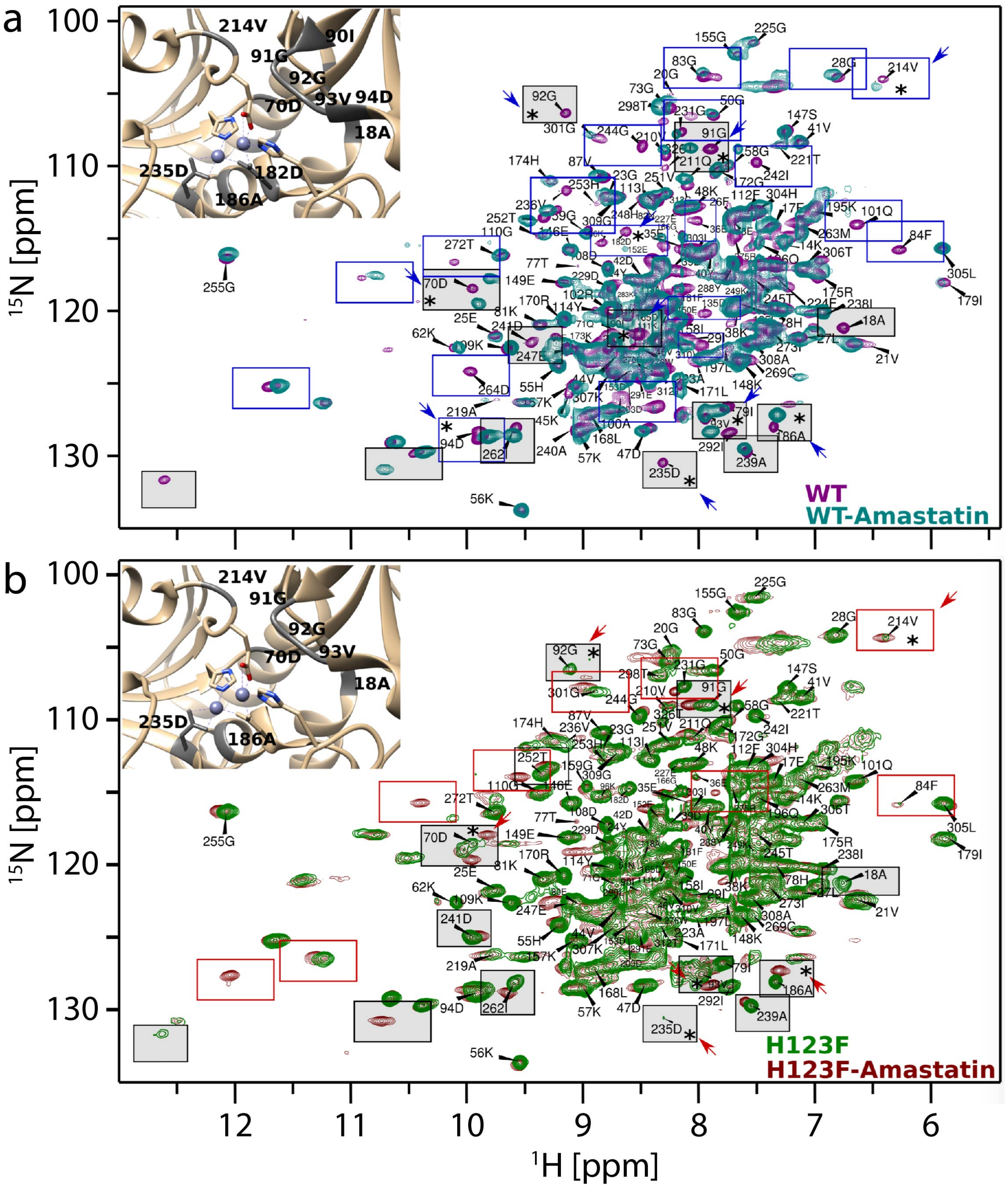

**Fig. S15.** Amastatin binding to wild-type (a) and H123F (b) TET2, monitored by $^{1}H$-$^{15}N$ MAS NMR spectra. Peaks corresponding to residues with significant chemical-shift perturbations are highlighted.

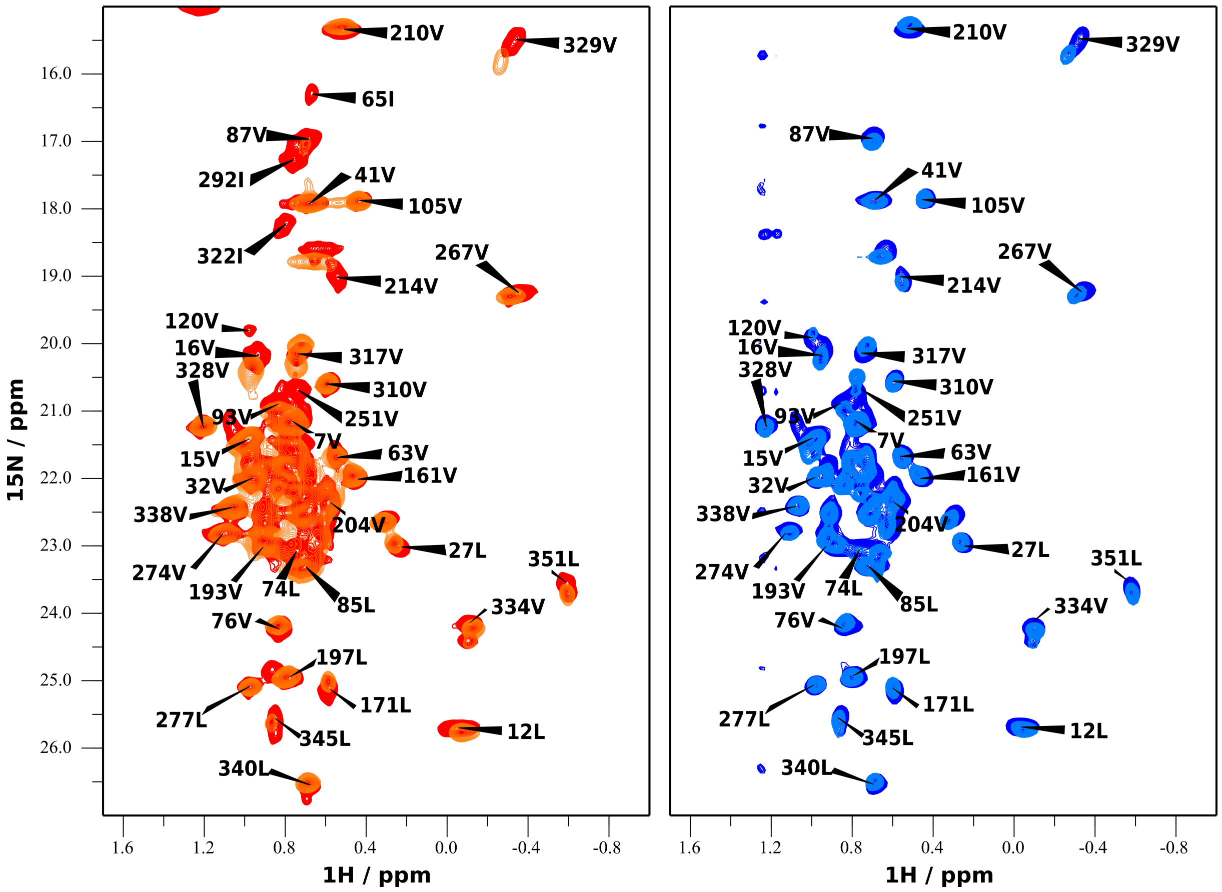


**Fig. S16.** Amastatin binding to wild-type wild-type (left) and H123F (right) TET2, monitored by $^{1}$H-$^{13}$C MAS NMR spectra.

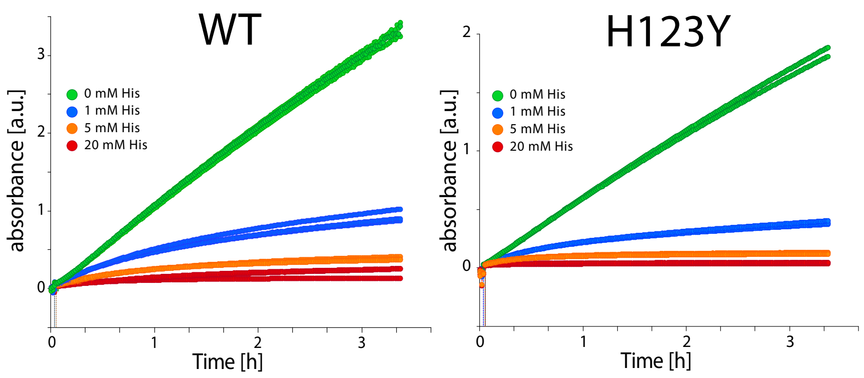


**Fig. S17.** Activity measurements in the presence of free histidine in solution, using the Leu-pNA assay. Shown are data for WT (left) and H123Y (right) TET2 in the presence of 0, 1, 5 and 20 mM histidine in the buffer. The experiment aims to see if a free histidine may have a (positive) effect on the catalytic activity, i.e. whether the histidine in the loop (H123) participates in the active site and if this function may be recapitulated by free histidine. If so, then the mutant that lacks the native H123 (right) may regain its activity in the presence of histidine. The opposite is observed, i.e. free histidine reduces the activity, and at 20 mM His the activity is reduced to almost zero. A plausible interpretation is that the free amino acid competes with the substrate for the active site.

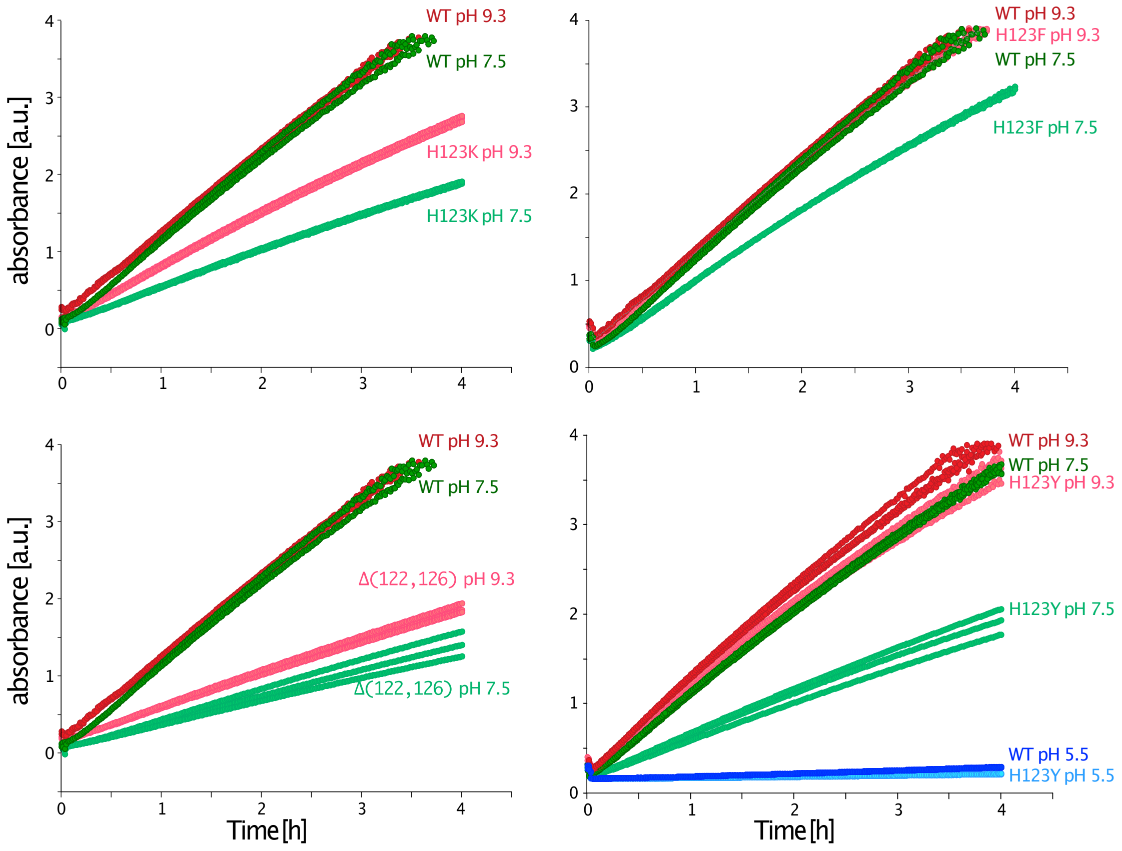


**Fig. S18.** Activity measurements of TET2 and loop-mutants at different pH values, using Leu-pNA as substrate. Shown are data for WT TET2 (displayed in all panels), H123K (top left), H123F (top right), the version with the shortened loop which retains the native His 123 (Δ(122,126), bottom left) and H123Y (bottom right).

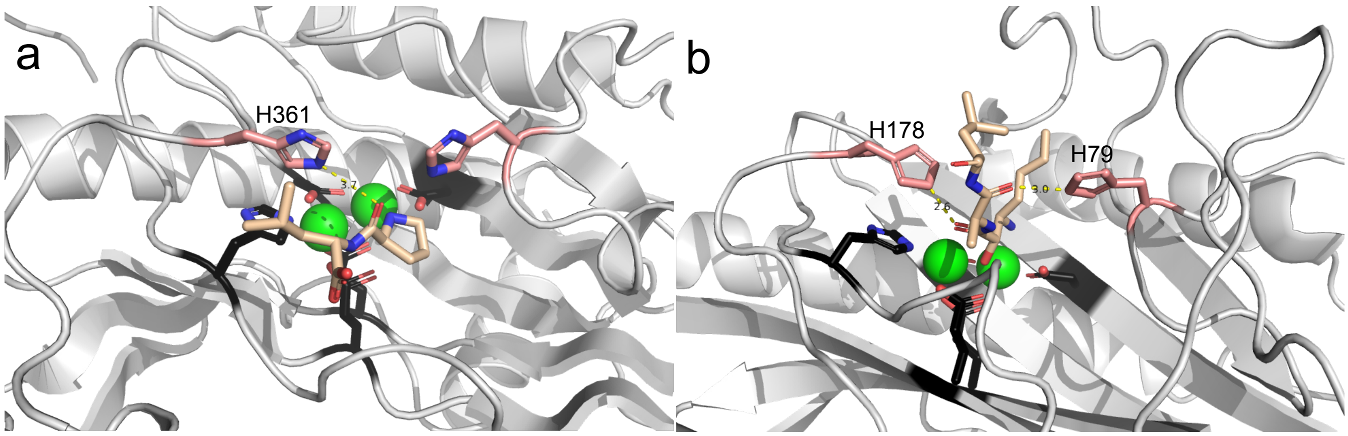


**Fig. S19.** Histidines in loop regions of the aminopeptidases AAP (panel a; PDB ID 1A16) and eMetAP (panel b; PDB ID 3MAT) interact with ligands in the active sites. In (a), a Pro-Leu is bound to a manganese form of aminopeptidase P (APP), directly above the water/hydroxide molecule that bridges between the metal ions. This state is thought to correspond to the binding mode of the C-terminal product of the hydrolysis reaction. H361 makes a water-mediated contact to an oxygen of the bound ligand (65). In (b), the inhibitor SL648, an analog of bestatin, is bound to *E. coli* methionine aminopeptidase, and the trapped states is thought to correspond to the transition state of the reaction. Two histidines make interactions with oxygens of the inhibitor, via the nitrogen of the imidazolium side chain, H79 and H178 (corresponding to H361 of APP) (66).

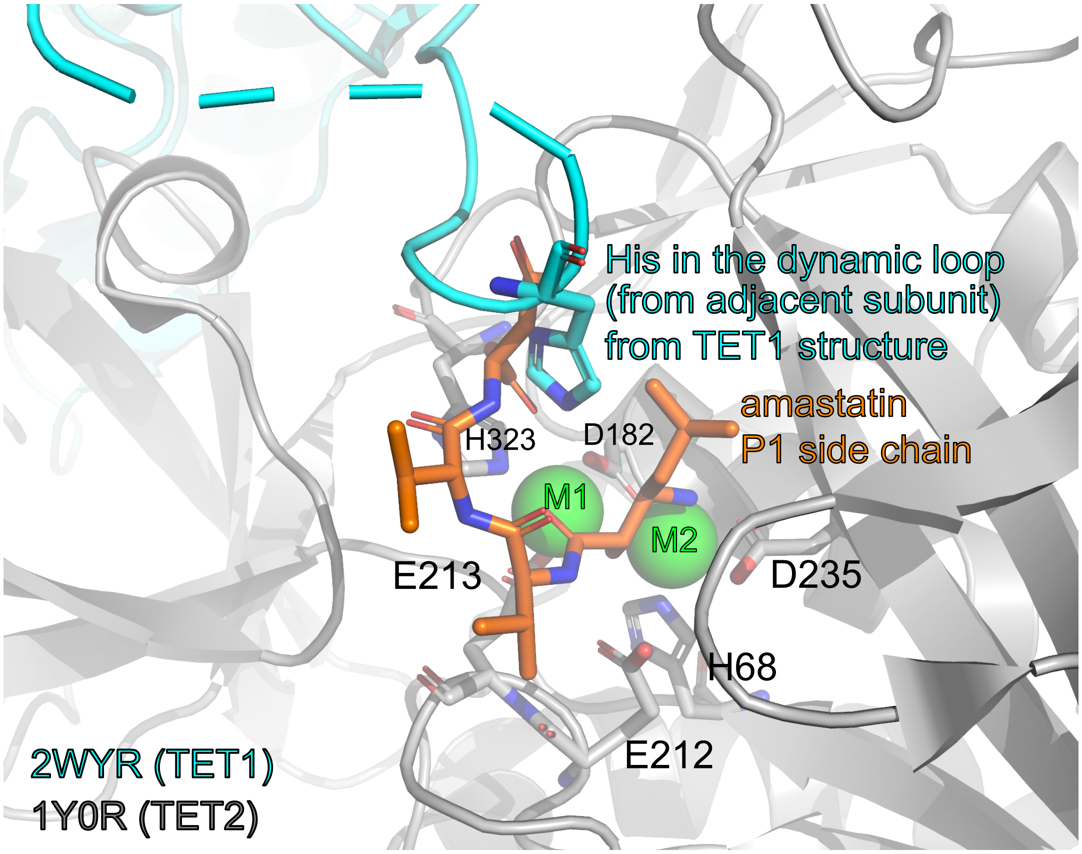


**Fig. S20.** Plausible model of how a substrate bound to the active site of a TET subunit might be approached by the histidine of the flexible loop of the adjacent subunit. Shown is the crystal structure of TET2 in complex with amastatin. Superimposed onto the structure of the adjacent subunit is the crystal structure of TET1, for which at least part of the loop has been modeled in the X-ray diffraction data. (Dashes indicate the part lacking in the structure 2WYR.) Note that among the known structure in which the His is modeled, this one has the His closest to the active site (cf. Fig. S13). It should be noted that in this rough model, multiple clashes would occur between the flexible loop and the substrate. However, given the significant degree of flexibility of both the loop and the substrate (at least beyond the first two residues), those may readily be avoided in such a complex. The loop of TET2 is approximately the same length as in TET1 (res. 119/133/H123 in TET2 correspond to res. 111/124/H115 in TET1. Thus, the His in TET2 has a similar "reach" towards the substrate as shown here. Given the uncertainty of the positioning of the His (Fig. S13), we refrain from proposing an exact pose of the side chain and mechanism of substrate-interaction.

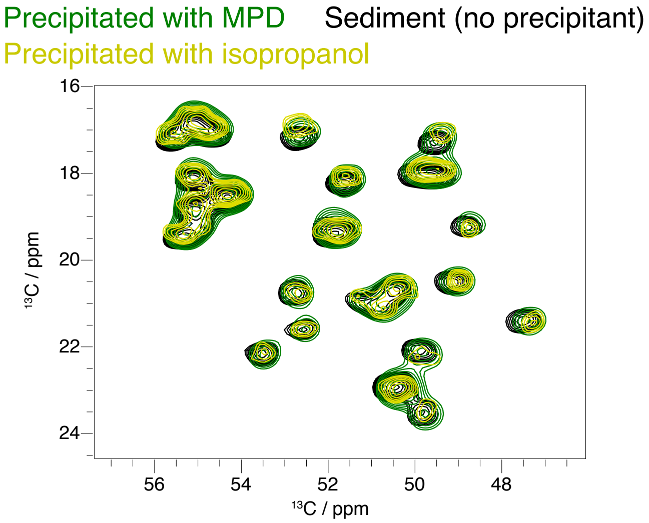


**Fig. S21.** Different preparations of TET2 MAS-NMR samples yield very similar spectra, indicating that the structure and dynamics are not strongly dependent on the sample preparation. All spectra are extracts from 2D carbon-carbon correlation spectra from MAS NMR (16.5 kHz MAS, 600 MHz $^{1}$H Larmor frequency, $^{13}$C,$^{15}$N-labeled samples, sample temperature ca. 25 $^{\circ}$C, DREAM mixing).
